# SGC-CLK-1 (CAF-170) a chemical probe for the Cdc2-Like kinases CLK1, CLK2, and CLK4

**DOI:** 10.1101/2022.12.15.520623

**Authors:** Deanna Tiek, Carrow I. Wells, Martin Schröder, Xiao Song, Carla Alamillo-Ferrer, Anshika Goenka, Rebeca Iglesia, Minghui Lu, Bo Hu, Frank Kwarcinski, Parvathi Sintha, Chandi de Silva, Mohammad Anwar Hossain, Alfredo Picado, William Zuercher, Reena Zutshi, Stefan Knapp, Rebecca B. Riggins, Shi-Yuan Cheng, David H. Drewry

## Abstract

Small molecule modulators are important tools to study both basic biology and the complex signaling of protein kinases. The cdc2-like kinases (CLK) are a family of four kinases that have garnered recent interest for their involvement in a diverse set of diseases such as neurodegeneration, autoimmunity, and many cancers. Targeted medicinal chemistry around a CLK inhibitor hit identified through screening of a kinase inhibitor set against a large panel of kinases allowed us to identify a potent and selective inhibitor of CLK1, 2 and 4. Here, we present the synthesis, selectivity, and potential binding site of this compound – SGC-CLK-1. We further show CLK2 has the highest binding affinity, and high CLK2 expression correlates with a lower IC_50_ in a screen of multiple cancer cell lines. Finally, we show that SGC-CLK-1 not only reduces serine arginine rich (SR) protein phosphorylation, but also alters SR protein and CLK2 subcellular localization in a reversible way. Therefore, we anticipate that this compound will be a valuable tool for increasing our understanding of CLKs and their targets, SR proteins, at the level of phosphorylation and subcellular localization.

## 1. Introduction

Kinases are attractive targets for drug discovery, with well documented success stories over the past two decades, and to date over 65 kinase inhibitors have been approved for clinical use by the FDA [1, 2]. The utility of kinase inhibitors stems from their intimate involvement in most physiological processes and pathways in healthy cells and tissues, and aberrant signaling through these same kinases, brought about by mutation or overexpression, for example, can lead to disease. One such important biological process regulated by the activity of kinases is pre-mRNA splicing, the process by which pre-mRNAs are processed into mature mRNAs that will be translated into proteins. The splicing process allows for the generation of alternately spliced mRNA variants, leading to the production of different protein products from the same gene [3, 4].

Recently, modulators of splicing, including intervention through so-called splicing kinases, have garnered attention for their therapeutic potential [5–7]. Splicing modulators have been suggested as potential therapeutics for human diseases including autoimmune diseases like lupus and rheumatoid arthritis [8], muscle diseases such as Duchenne muscular dystrophy [9, 10], neurodegenerative diseases like Alzheimer’s diseases and Parkinson’s disease [11], and various types of cancers [12–20]. Alternate splicing influences numerous cellular processes that are considered to be hallmarks of cancer [21–23], and the body of evidence for the role of the splicing process in tumorigenesis has led to the suggestion of including aberrant alternative splicing as an additional hallmark of cancer [24].

The cdc2-like kinases – CLK1, CLK2, CLK3, and CLK4 – constitute a small family of kinases that are part of the CMGC group (CDK, MAPK, GSK3, and CLK) of the human kinome, and they play key roles in the splicing process. CLKs can phosphorylate (p) serine-arginine rich (SR) proteins, and this phosphorylation re-localizes the pSR proteins from their holding area in the nuclear speckles to the spliceosome to dictate exon inclusion/exclusion in the nascent transcript [25–27]. As expected, the four CLK family members are highly homologous, but there are differences in sequence that give rise to functional differences and lead to different compound inhibition profiles across the four CLKs [7,14,28,29].

As targeting splicing through splicing modulators has becomes an active and attractive area of research, a growing number of small molecule inhibitors have been created to better understand the CLKs in physiological roles such as splicing, and to investigate the consequences of CLK inhibition in disease-relevant models. CLK inhibitors and the therapeutic potential of CLK inhibition have been reviewed recently [7,14,29]. Several CLK inhibitors have been sufficiently characterized so that they can be considered chemical probes for the CLKs. **Table 1** depicts the recent CLK inhibitors that can be used to probe CLK biology, along with examples of CLK inhibitors that have advanced into clinical trials.

**Table 1:**
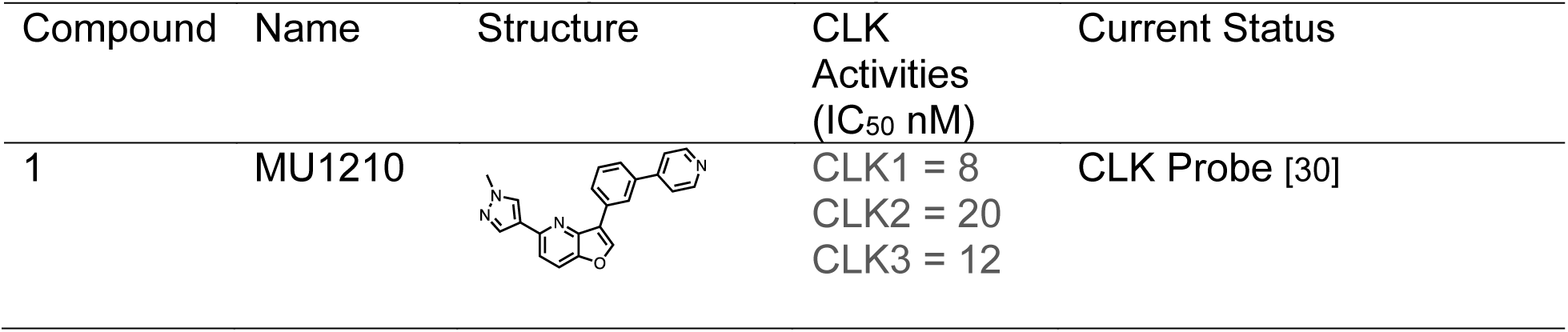
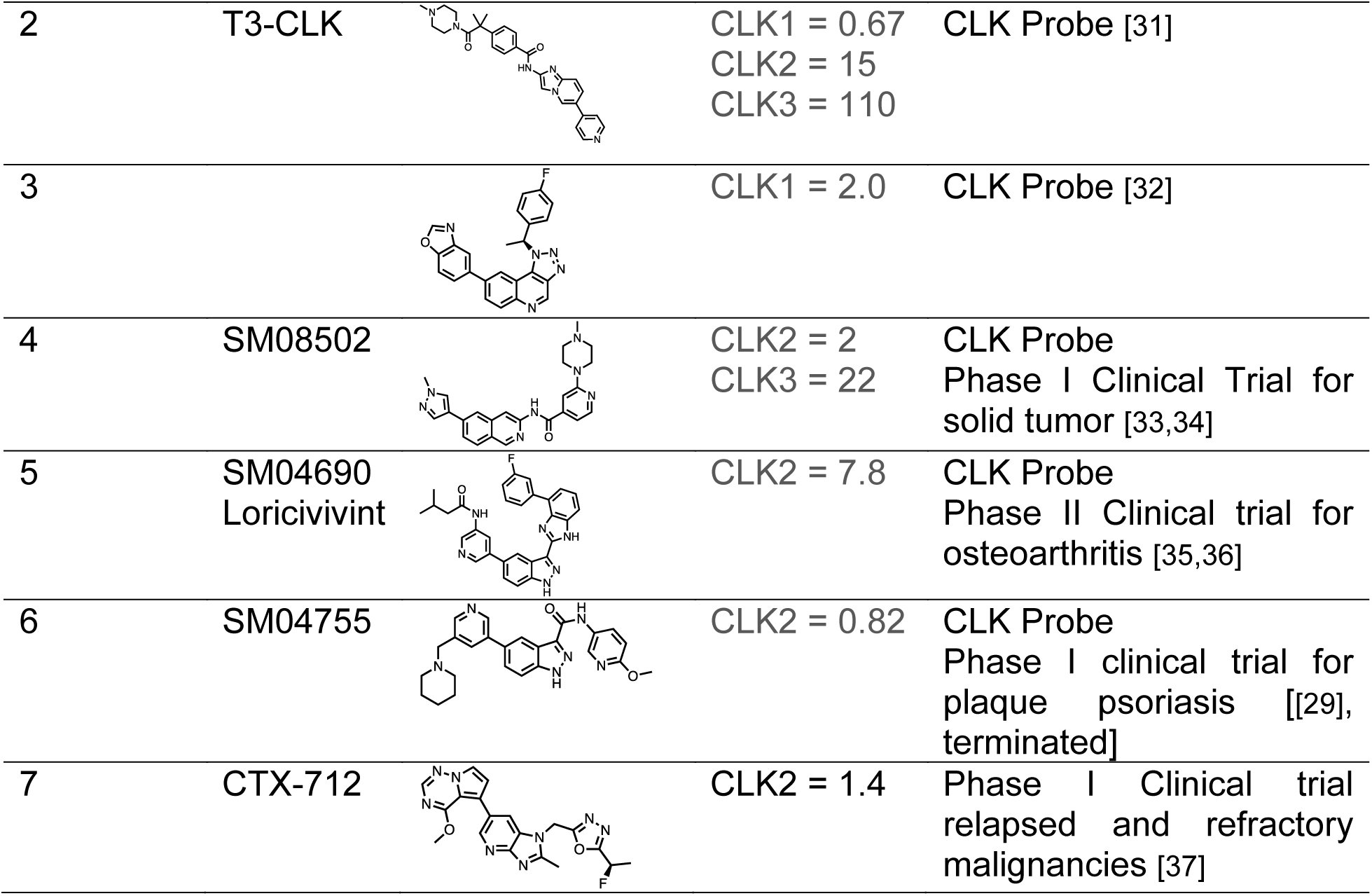
literature CLK chemical probes and compounds in clinical trials

| Compound | Name | Structure | CLK Activities<br>(IC<sub>50</sub> nM) | Current Status |
| --- | --- | --- | --- | --- |
| 1 | MU1210 |  | CLK1 = 8<br>CLK2 = 20<br>CLK3 = 12 | CLK Probe [30] |
| 2 | T3-CLK |  | CLK1 = 0.67<br>CLK2 = 15<br>CLK3 = 110 | CLK Probe [31] |
| 3 |  |  | CLK1 = 2.0 | CLK Probe [32] |
| 4 | SM08502 |  | CLK2 = 2<br>CLK3 = 22 | CLK Probe<br>Phase I Clinical Trial for solid tumor [33,34] |
| 5 | SM04690<br>Loricivivint |  | CLK2 = 7.8 | CLK Probe<br>Phase II Clinical trial for osteoarthritis [35,36] |
| 6 | SM04755 |  | CLK2 = 0.82 | CLK Probe<br>Phase I clinical trial for plaque psoriasis [[29], terminated] |
| 7 | CTX-712 |  | CLK2 = 1.4 | Phase I Clinical trial relapsed and refractory malignancies [37] |

From the recent CLK reviews and the primary CLK medicinal chemistry publications it becomes apparent that CLK inhibitor scaffolds have varied potency across the CLK family and carry different off-target kinase activities. There are also examples of kinase inhibitors initially developed for other enzymes that can act as CLK inhibitors [38–41]. As the CLKs share very similar ATP binding pockets, creating a potent and selective tool molecule for each CLK has proven difficult to date. Nevertheless, there are several ongoing clinical trials currently underway with CLK inhibitors ranging from osteoporosis to neurodegenerative disorders and cancer (**Supplemental Table 1**). Alternate scaffolds with different pharmacological profiles, intrafamily potency profiles, and kinase off-target profiles should prove useful in the quest to find useful CLK inhibitors that can modulate disease phenotypes and serve as precursors for new therapeutics.

Here, we report a new potent and cell active CLK chemical probe, CAF-170, renamed as SGC-CLK-1, and a negative control compound CAF-225, renamed as SGC-CLK-1N. SGC-CLK-1 (CAF-170) has excellent kinome wide selectivity, is a potent inhibitor of CLK1, CLK2, and CLK4, and produces a unique phenotype outside of the normal presumed CLK inhibitor function of simply decreasing SR phosphorylation. At low nanomolar concentrations, we observe that SGC-CLK-1 re-distributes CLK2 and pSRs leading to a decrease in growth of both melanoma and glioblastoma cells. Finally, this relocalization is reversible, which positions SGC-CLK-1 as a useful mechanistic tool for CLK2 function and a scaffold for further optimization.

## 2. Materials and methods

### 2.1 Cell lines and culturing conditions

MDA-MB-435 (ATCC) and U118-MG (ATCC) were cultured in DMEM (Thermo Fisher Scientific, 11995-065) and supplemented with 10% FBS (Thermo Fisher Scientific, 10437028). All cells were authenticated by short tandem repeat analysis at IDEXX BioAnalytics, and were cultured for less than 10 passages prior to use.

### 2.2 Western blotting

Cells were lysed in RIPA buffer supplemented with protease and phosphatase inhibitors (Roche (Sigma), 4906837001) for protein extractions and separated by polyacrylamide gel electrophoresis using 4-12% gradient gels (Novex by Life Tech, NP0321BOX). They were then transferred onto Nitrocellulose membranes (Invitrogen, IB23001) with the iBlot2 (Invitrogen, IB21001) and probed with the following antibodies post 1 hr blocking in 5% milk: anti-phosphorylated (p)SR proteins (clone 1H4, 1:500, MABE50; MilliporeSigma), and anti–*α/β*-tubulin (1:1000, Cell Signaling Technology, 2148) antibodies. Following washing with TBS-T, the blot was incubated with corresponding HRP-conjugated secondary antibodies (DAKO, anti-rabbit immunoglobulins, P0217; anti-mouse immunoglobulins, P0260). Blots were developed with enhanced chemiluminescence

### 2.3 Cell proliferation assays

Cells were seeded in 96-well plastic tissue culture plates at 1000 cells/well 1 d prior to treatment with the indicated concentrations of CAF-170 or the negative control compound CAF-225. Cells were treated for a total of 8 days, with medium changed and drug replenished on day 4. Staining with crystal violet, re-solubilization, and analysis of staining intensity as a proxy for cell number was performed as previously described [42].

### 2.4 Immunofluorescence

Cells were seeded at a density of 45,000–55,000 cells onto 18-mm-diameter #1.0 round coverslips (VWR, Radnor, PA, USA) in 12-well dishes. On the following day, the cells were treated with indicated concentrations of inhibitor (500 nM CAF-225, 100nM, 500nM CAF-170, 500 nM MU140, 500 nM MU1210, 500 nM T3) for either 15’, 30’, 45’, or 60’ as indicated in the figure legends in conditioned media from cells seeded to the same density. Media was removed, and coverslips were washed 3X with PBS and then fixed and permeabilized in 3.2% paraformaldehyde with 0.2% Triton X-100 in PBS for 5 min at room temperature. Three washes were performed with PBS in the 12-well plate, and then coverslips were inverted onto 120 μl of primary antibody in the antibody block (0.1% gelatin with 10% normal donkey serum in PBS) on strips of parafilm and incubated for 1 hr. Coverslips were first incubated with anti-pSR (1:150) and anti-CLK2 (1:500, Sigma Aldrich, HPA055366-100UL) antibodies for 1 hr. After incubation with primary antibodies, coverslips were washed 3 times with PBS. Then, coverslips were inverted onto 100 μl of antibody block with secondary antibodies (Alexa Fluor 488 anti-mouse, 1:200, A11029; Thermo Fisher Scientific) and DAPI (DNA, 1:500 dilution) for 20 min in the dark. Coverslips were again washed 3 times with PBS and then gently dipped 4 times into molecular biology-grade water before inversion onto 1 drop of Fluoro-Gel (with N-[tris(hydroxymethyl)methyl]-2-aminoethanesulfonic acid buffer, 17985–30; Electron Microscopy Sciences, Hatfield, PA, USA) and allowed to air-dry in the dark for at least 10 min. Slides were stored at 4°C until image collection on an Olympus BX-53 microscope with a series 120 Q X-cite Lumen Dynamics laser.

### 2.5 General chemistry and compound synthesis information

Reagents were obtained from reputable commercial vendors. Solvent was removed via rotary evaporator under reduced pressure. Thin layer chromatography was used to track reaction progress. These abbreviations are used in experimental procedures: mmol (millimoles), μmol (micromoles), mg (milligrams), mL (milliliters) and r.t. (room temperature). ^1^H NMR and/or additional microanalytical data was collected to confirm identity and assess purity of final compounds. Magnet strength for NMR spectra is included in line listings. Peak positions are listed in parts per million (ppm) and calibrated versus the shift of CD3OD-d_4_; coupling constants (*J* values) are reported in hertz (Hz); and multiplicities are as follows: singlet (s), doublet (d), doublet of doublets/triplets (dd/dt), doublet of doublets of triplets (ddt), triplet (t), and multiplet (m). Compounds were confirmed to be >95% pure by ^1^H NMR and HPLC analysis.

### 2.6 Probe and negative control synthesis

The synthetic route to make the chemical probe SGC-CLK-1 is depicted in Scheme 1.

**Scheme 1:**
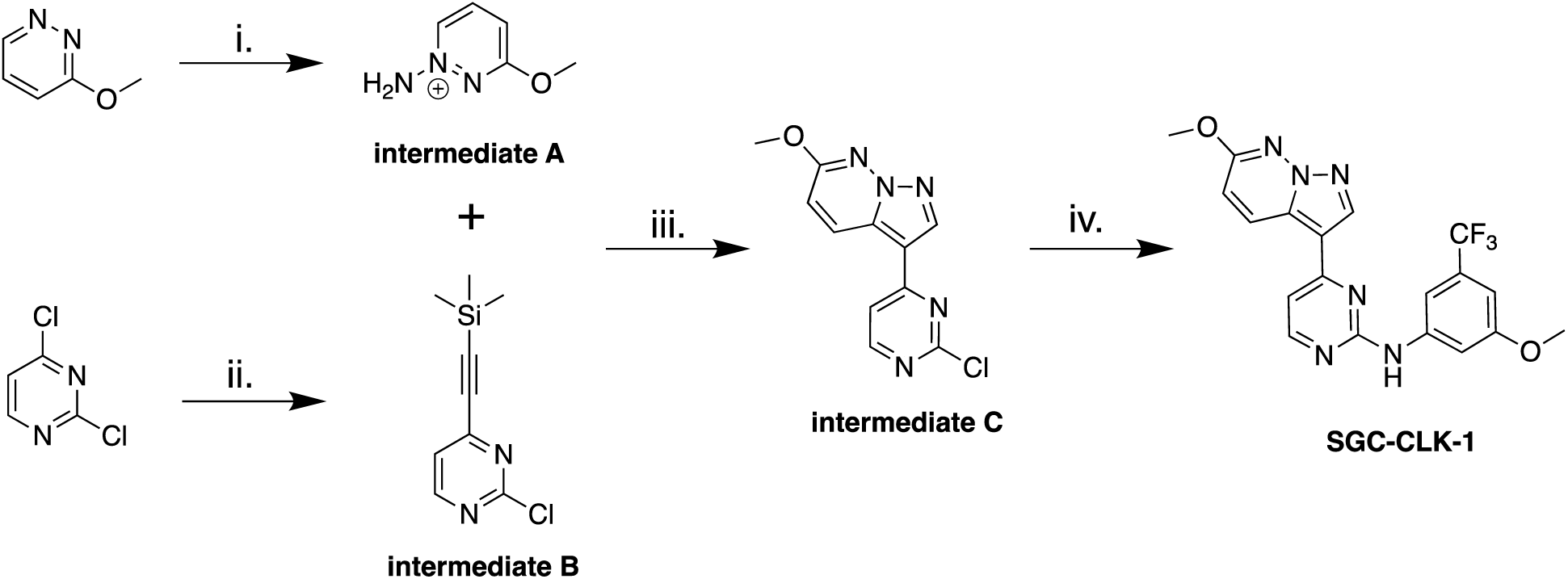
Chemistry route for probe synthesis. Reagent and conditions: i) hydroxylamine-O-sulfonic acid (HOSA), KHCO_3_, H_2_O, 80 C, 14 h; ii) KHCO_3_, KOH, H_2_O, DCM, r.t, 18 h; iii) ethynyltrimethyl silane, Pd(dppf)Cl_2_ · CH_2_Cl_2_, CuI, PPh_3_, TEA, THF, 70 C, 15 min; iv) TFA, tert-BuOH, 85 C, 15 h.

#### Synthesis of Intermediate A: 1-amino-3-methoxypyridazin-1-ium

Hydroxylamine-O-sulfonic acid (HOSA) (0.83 g, 7.39 mmol, 3.7 eq.) was dissolved in water (1.6 mL), and then treated with KHCO_3_ (0.74 g, 7.39 mmol, 3.7 eq.) in water (1.0 mL), pH 5. Then 3-methoxypyridazine (0.22 g, 2.00 mmol, 1.00 eq.) was added in portions. The reaction was then stirred at 80 °C overnight. This solution of the crude material was used for the next reaction without purification.

#### Synthesis of Intermediate B: 2-chloro-4-((trimethylsilyl)ethynyl)pyrimidine

2,4-dichloropyrimidine (1.00 g, 6.7 mmol, 1 eq.) in tetrahydrofuran (30 mL) was degassed with nitrogen for 10-12 min, then ethynyltrimethylsilane (0.73 g, 7.4 mmol, 1.1 eq.) and triethylamine (0.74 g, 7.5 mmol, 1.1 eq.) were added, and the reaction degassed for another 5min. At this time PdCl2(dppf)-CH2Cl2adduct (0.27 g, 0.34 mmol, 0.05 eq.), copper(I) iodide (0.13 g, 0.67 mmol, 0.10 eq.), and triphenylphosphine (0.18 g, 0.67 mmol, 0.10 eq.) were added. The reaction was then refluxed for 15 min and copious solids formed. By TLC, all starting material was consumed (Hexanes:EtOAc, 85:15). Hexanes was added to precipitate triphenylphosphine oxide, and the reaction was filtered and rinsed with EtOAc. The crude material was concentrated in the rotavap, to provide a thick brown liquid. The desired product was obtained by flash chromatography using EtOAc in hexanes (0% to 20%). The desired fractions were concentrated in the rotary evaporator, dried under vacuum overnight, to provide the desired product as a brown solid 770 mg (yield 50%), purity>90% by NMR.

^1^H NMR (400 MHz, DMSO-*d*_6_) δ ppm 0.27 (s, 9 H), 7.66 (d, *J*=5.1 Hz, 1 H), 8.80 (d, *J*=5.1 Hz, 1 H). ^13^C NMR (101 MHz, DMSO-*d*_6_) δ ppm 100.4, 102.1, 122.6, 151.4, 160.1, 161.2.

#### Synthesis of Intermediate C: 3-(2-chloropyrimidin-4-yl)-6-methoxypyrazolo[1,5-b]pyridazine

The aqueous solution of 1-amino-3-methoxypyridazin-1-ium (0.20 g, 1.6 mmol, 1.5 eq.) from the previous step (pH = 1) was treated with saturated KHCO_3_ to bring the pH to 7. 2-chloro-4-((trimethylsilyl)ethynyl)pyrimidine (0.22 g, 1.04 mmol, 1.00 eq.) was dissolved in 1 mL of DCM (1 M), and added in one portion to the crude aminated pyridazine. KOH (0. 320 g, 6.26 mmol, 6 eq.) was dissolved in H_2_O (5 mL) 1.0 M and was added in one portion to the reaction mixture. The reaction mixture turned dark red in color after 5-10 min. The reaction mixture was stirred vigorously at r.t. for 22 hr. The crude mixture was then quenched with water, extracted with DCM, and the combined organic layers were dried over anhydrous Na_2_SO_4_. Filtration and rotary evaporation of the solvent provide the crude product which was dry loaded on a 10 g Biotage Sfar 60 µm silica cartridge (Hexanes/EtOAc 70:30) and chromatographed to provide a light pink solid (0.095 g, 35% yield). LCMS [M+1] = 262, purity > 95%.

^1^H NMR (400 MHz, DMSO-*d*_6_) δ ppm 4.02 (s, 3 H), 7.27 (d, *J*=9.4 Hz, 1 H), 8.01 (d, *J*=5.5 Hz, 1 H), 8.66 - 8.78 (m, 2 H), 8.83 (s, 1 H).

#### Synthesis of SGC-CLK-1 (CAF-170): N-(3-methoxy-5-(trifluoromethyl)phenyl)-4-(6-methoxypyrazolo[1,5-b]pyridazin-3-yl)pyrimidin-2-amine

3-(2-chloropyrimidin-4-yl)-6-methoxypyrazolo[1,5-b]pyridazine (0.085 g, 0.32 mmol, 1 eq.), 3-methoxy-5-(trifluoromethyl)aniline (0.075 g, 0.39 mmol, 1.20 eq.), and tert-butanol (4.5 mL) were combined in a microwave vial, 4 small drops of TFA added, the vial was sealed, and the reaction stirred at 85 C for 15 hr. The reaction mixture was allowed to come to r.t. and quenched with water. Using aqueous NaHCO_3_, the pH was adjusted to 7 and a solid precipitated. More water was added, the solid was obtained by filtration, thoroughly rinsed with water, and air dried. A pale pink solid was obtained, 110 mg recovered. The product was purified using Biotage Sfar 10g silica cartridge, solid load and a Hexanes/EtOAc gradient from 0% to 50% EtOAc to provide the desired product as a pale off-white solid (0.057 g, > 95% pure). LCMS [M+1] = 417, 418. 1H NMR (850 MHz, DMSO-d_6_) δ 9.90 (s, 1H), 9.02 (d, J = 9.6 Hz, 1H), 8.72 (s, 1H), 8.54 (d, J = 5.2 Hz, 1H), 7.84 (d, J = 1.8 Hz, 1H), 7.68 (t, J = 2.2 Hz, 1H), 7.43 (d, J = 5.2 Hz, 1H), 7.15 (d, J = 9.5 Hz, 1H), 6.85 (t, J = 2.0 Hz, 1H), 4.03 (s, 3H), 3.84 (s, 3H). ^13^C NMR (214 MHz, DMSO-d_6_) δ 163.22, 162.93, 162.78, 162.58, 161.21, 145.80, 141.40, 133.84, 133.54, 133.39, 132.60, 127.97, 126.70, 116.29, 113.54, 111.55, 111.00, 110.54, 106.06, 58.69, 57.91.

#### The synthetic route to make the negative control compound SGC-CLK-1N is depicted in Scheme 2

**Scheme 2:**
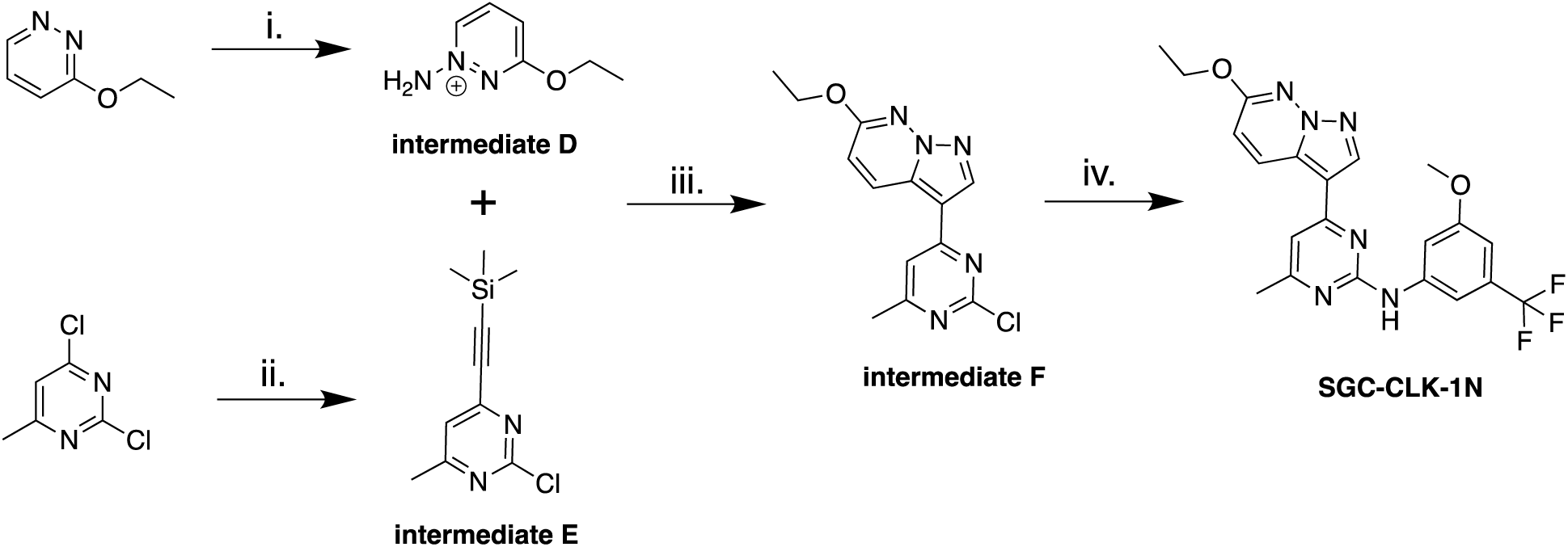
Synthesis of negative control compound SGC-CLK-1N. Reagents and conditions: i) hydroxylamine-O-sulfonic acid (HOSA), KHCO_3_, H_2_O, 80 C, 14 h; ii) KHCO_3_, KOH, H_2_O, DCM, r.t, 18 h; iii) ethynyltrimethylsilane, Pd(dppf)Cl_2_ · CH_2_Cl_2_, CuI, PPh_3_, TEA, THF, 70 C, 15 min; iv) TFA, tert-BuOH, 85 °C, 15 hr.

#### Synthesis of Intermediate D. 1-amino-3-ethoxypyridazin-1-ium

Hydroxylamine-O-sulfonic acid (HOSA), (0.82 g, 7.25 mmol, 4.5 eq.) was dissolved in water (1.6 mL), then treated with KHCO_3_ (0.72 g, 7.25 mmol, 4.5 eq.) in water (1.0 mL), pH 5. 3-ethoxypyridazine (0.20 g, 1.61 mmol, 1.00 eq.) was added to the HOSA solution portionwise. The reaction was then stirred at 80 °C overnight. The resulting crude solution was used in the next reaction without purification.

#### Synthesis of Intermediate E. 2-chloro-4-methyl-6-((trimethylsilyl)ethynyl) pyrimidine

A solution of 2,4-dichloro-6-methylpyrimidine (1.0 g, 6.10 mmol, 1.00 eq.) dissolved in THF (30 mL) was degassed with nitrogen for 10-12 min, then ethynyltrimethylsilane (0.66 g, 6.7 mmol, 1.1 eq.) and triethylamine (0.68 g, 6.7 mmol, 1.10 eq., 0.94 mL) were added and the reaction mixture was degassed for another 5 min, and then PdCl2(dppf)-CH_2_Cl_2_adduct (0.25 g, 0.31 mmol, 0.05 eq.), copper(I) iodide (0.12 g, 0.61 mmol, 0.10 eq.), and triphenylphosphine (0.16 g, 0.61 mmol, 0.10 eq.) were added. The reaction mixture was refluxed for 15 min and a precipitate formed. The crude reaction mixture was filtered over celite, and then the celite plug rinsed with EtOAc. The EtOAc washes were concentrated by rotary evaporation to provide a thick brown oil. The crude product was purified using flash chromatography (0 to 15% EtOAc) to provide 1.03 g of the desired product (yield 67%), purity around 90%.

^1^H NMR (400 MHz, DMSO-*d*_6_) δ ppm 0.27 (bs, 9 H), 2.46 (s, 3 H), 7.61 (s, 1 H).

#### Synthesis of Intermediate F. 3-(2-chloro-6-methylpyrimidin-4-yl)-6-ethoxypyrazolo[1,5-b]pyridazine

The crude solution of 1-amino-3-ethoxypyridazin-1-ium (0.22 g, 1.6 mmol, 2.0 eq.) from the previous step (pH = 1) was treated with saturated KHCO_3_ to bring the pH to 7. 2-chloro-4-methyl-6-((trimethylsilyl)ethynyl)pyrimidine (0.18 g, 0.88 mmol, 1.00 eq.) was dissolved in 0.8 mL of DCM (1 M), and added in one portion to the crude aminated pyridazine. KOH (0. 27 g, 4.8 mmol, 6 eq.) dissolved in H_2_O (4.8 mL) 0.9-1.0 M was then added in one portion to the reaction mixture. The reaction mixture turned dark red in color after 5-10 min. The reaction mixture was stirred vigorously at r.t. for 22 hr. The crude mixture was then quenched with water, extracted with DCM, and the combined organic layers were dried over anhydrous Na_2_SO_4_. Filtration and evaporation of the solvent proved the crude product which was dry loaded on a 10 g Biotage Sfar 60 um silica cartridge. Chromatography using Hexanes/EtOAc 7(0:30) provided the desired product as an off-white solid (0.116 g, 48% yield). LCMS [M+1] = 290, purity > 95%.

^1^H NMR (400 MHz, DMSO-*d*_6_) δ ppm 1.41 (t, *J*=7.0 Hz, 3 H), 2.49 (br s, 3 H), 4.41 (q, *J*=7.0 Hz, 2 H), 7.23 (d, *J*=9.4 Hz, 1 H), 7.92 (s, 1 H), 8.69 - 8.77 (m, 2 H).

#### Synthesis of Negative control SGC-CKL-1N. 4-(6-ethoxypyrazolo[1,5-*b*]pyridazin-3-yl)-*N*-(3-methoxy-5-(trifluoromethyl)phenyl)-6-methylpyrimidin-2-amine

3-(2-chloro-6-methylpyrimidin-4-yl)-6-ethoxypyrazolo[1,5-b]pyridazine (0.065 g, 0.22 mmol, 1 eq.), 3-methoxy-5-(trifluoromethyl)aniline (0.051 g, 0.27 mmol, 1.20 eq.), and tert-butanol (4.5 mL) were mixed into a microwave vial, 4 small drops of TFA added, the vial was sealed, and the reaction stirred at 85 °C for 15 hr. The reaction mixture was allowed to cool to r.t., and quenched with water. The pH was adjusted to 7 with aqueous NaHCO_3_, and a solid precipitated. More water was added, and the resulting solid was filtered, thoroughly rinsed with water, and air dried to provide a pale pink solid (110 mg recovered). The product was purified using Biotage Sfar 10 g silica cartridge, solid load, eluting with Hexanes/EtOAc gradient from 0% to 50% EtOAc to provide the desired product as an off-white solid (0.065 g, yield 30%, > 95% pure). LCMS [M+1] = 445, 446.

1H NMR (850 MHz, DMSO-d_6_) δ 9.85 (s, 1H), 8.98 (d, J = 9.3 Hz, 1H), 8.65 (s, 1H), 7.84 (d, J = 1.8 Hz, 1H), 7.75 (s, 1H), 7.35 (s, 1H), 7.09 (d, J = 9.5 Hz, 1H), 6.83 (t, J = 2.0 Hz, 1H), 4.42 (q, J = 7.0 Hz, 2H), 3.84 (s, 3H), 2.44 (s, 3H), 1.43 (t, J = 7.0 Hz, 3H). _13_C NMR (214 MHz, DMSO-d_6_) δ 170.71, 163.20, 162.67, 162.40, 162.26, 145.99, 141.02, 133.78, 133.48, 133.33, 132.48, 127.99, 126.71, 116.20, 113.69, 110.82, 110.65, 110.48, 105.90, 66.43, 58.62, 27.01, 17.24.

### 2.7 Kinome-wide selectivity analysis and confirmation of binding

The scanMAX assay platform offered by the Eurofins DiscoverX Corporation was used to assess the selectivity of the chemical probe when screened at 1 µM. This platform measures the binding of a compound to 403 wildtype (WT) human as well as several mutant and non-human kinases, generating percent of control (PoC) values for every kinase evaluated (Davis et al., 2011). A selectivity score (S_10_(1 µM)) is calculated using the PoC values for WT human kinases only.

### 2.8 NanoBRET assays

The CLK family of nanoBRET assays were run as directed by Promega. Briefly, to 10.5mL of 2.0 x 10^5 of Hek293 Cells (ATCC CRL1573) in growth media (DMEM + 10% FBS) was added 525uL of a 9:1 ratio of carrier DNA (Promega E4881) to a CLK nLuc construct-CLK1 CLK2 or CLK4 (Promega – NV1131, NV1141, NV1151) in OPTImem (Gibco) + 15.75uL of Fugene (Promega). To each well of a 96 well plate (Corning 3717) was then added 100uL of cells. The cells were then transfected overnight (20 hours) at 37degC. After overnight the media was aspirated and replaced with 85uL of OPTImem. To each well was then added 5uL of tracer K5 (Promega) at a final concentration of 0.5uM per well. Then 10uL of CLK inhibitor was added with concentrations varying from 1nM to 1uM. The plates were then returned to 37degC for 2 hours. After 2 hours the plates were equilibrated to room temperature for 15 minutes. After 15 minutes, 50uL of 3x complete substrate plus inhibitor solution (Promega) was added to each well and read after 2 min on a GloMax instrument. Test compounds were evaluated at eleven concentrations in competition with NanoBRET Tracer K5 in HEK293 cells transiently expressing the CLK(1,2, or 4) fusion protein. Raw milliBRET (mBRET) values were obtained by dividing the acceptor emission values (600 nm) by the donor emission values (450 nm) then multiplying by 1000. Averaged control values were used to represent complete inhibition (no tracer control: Opti-MEM + DMSO only) and no inhibition (tracer only control: no compound, Opti-MEM + DMSO + Tracer K5 only) and were plotted alongside the raw mBRET values. The data with n=3 biological replicates was first normalized and then fit using Sigmoidal, 4PL binding curve in Prism Software (version 8, GraphPad, La Jolla, CA, USA). All error bars are based on n=3 and are +/-standard error (SE).

### 2.9 CLK KinaseSeeker Assays

The Luceome KinaseSeeker™ assay [43, 44] was used to measure binding to CLK1, CLK2, and CLK3. Stock solutions (10 mM in DMSO) of test compounds were serially diluted in DMSO to make assay stocks. Prior to initiating screening or IC_50_ determination, the test compounds were evaluated for false positive results against split luciferase. The test compound was screened against CLK1, CLK2, and CLK3 at a minimum of eight different concentrations in duplicate. For kinase assays, a 24 mL aliquot of lysate containing Cfluc-kinase and Fos-Nfluc was incubated with either 1 μL of DMSO (for no-inhibitor control) or compound solution in DMSO for 2 h in the presence of a kinase-specific probe. Luciferin assay reagent (80 μL) was added to each solution, and luminescence was immediately measured on a luminometer. The % inhibition was calculated using the following equation: % inhibition = [ALU (control) − ALU (sample)]/ALU (control) × 100. For IC_50_ determinations, each compound was tested at a minimum of eight different concentrations. The % inhibition was plotted against compound concentration, and the IC_50_ value was determined for each compound using an eight-point curve.

## 3. Results

### Discovery and characterization of a CLK chemical probe

In our original kinase chemogenomic set known as PKIS (Published Kinase Inhibitor Set), we identified GW807982X (CAF-022) as a potent inhibitor of members of the CLK family with good kinome wide selectivity based on screening against a panel of over 200 kinases at NanoSyn. CAF-022 has subsequently been shown to selectively inhibit chemoresistant glioblastoma cells [20]. To further understand the selectivity of this compound we screened it at a concentration of 1 µM in the Eurofins KINOME*scan*® panel. In this assay format, GW807982X exhibited very good selectivity, with an S_10_ (1 µM) = 0.02, indicating that only 2% of the kinases tested had a PoC < 10 at a screening concentration of 1 µM. In this assay, lower PoC values arise from potent binding to the kinases in question. The kinome wide data is available as **Supplemental Table 1**. In efforts to further enhance selectivity, we made a series of analogs. We utilized the Luceome KinaseSeeker™ assays to assess the affinity towards CLK1, CLK2, and CLK3. Close analogue CAF-170 (**Figure 1a**) bound effectively to CLK1 with IC_50_ = 41 nM and CLK2 with IC_50_ = 36 nM, but when screened at a concentration of 1 µM against CLK3 showed no binding at all. CAF-170 demonstrated improved kinome-wide selectivity, with S_10_ (1 µM) = 0.002 in the KINOME*scan*® assay format (**Figure 1b** and **Supplemental Table 1**). CAF-170, renamed as SGC-CLK-1, bound to only six kinases in the KINOME*scan*® panel with a PoC<35: CLK1, CLK2, CLK4, HIPK1, HIPK2, and MAPK15 (also known as ERK8). Given this excellent selectivity we chose to characterize SGC-CLK-1 more fully in a series of orthogonal assays to assess its potential to serve as a cell active chemical probe for CLK1, CLK2, and CLK4.

**Figure 1:**
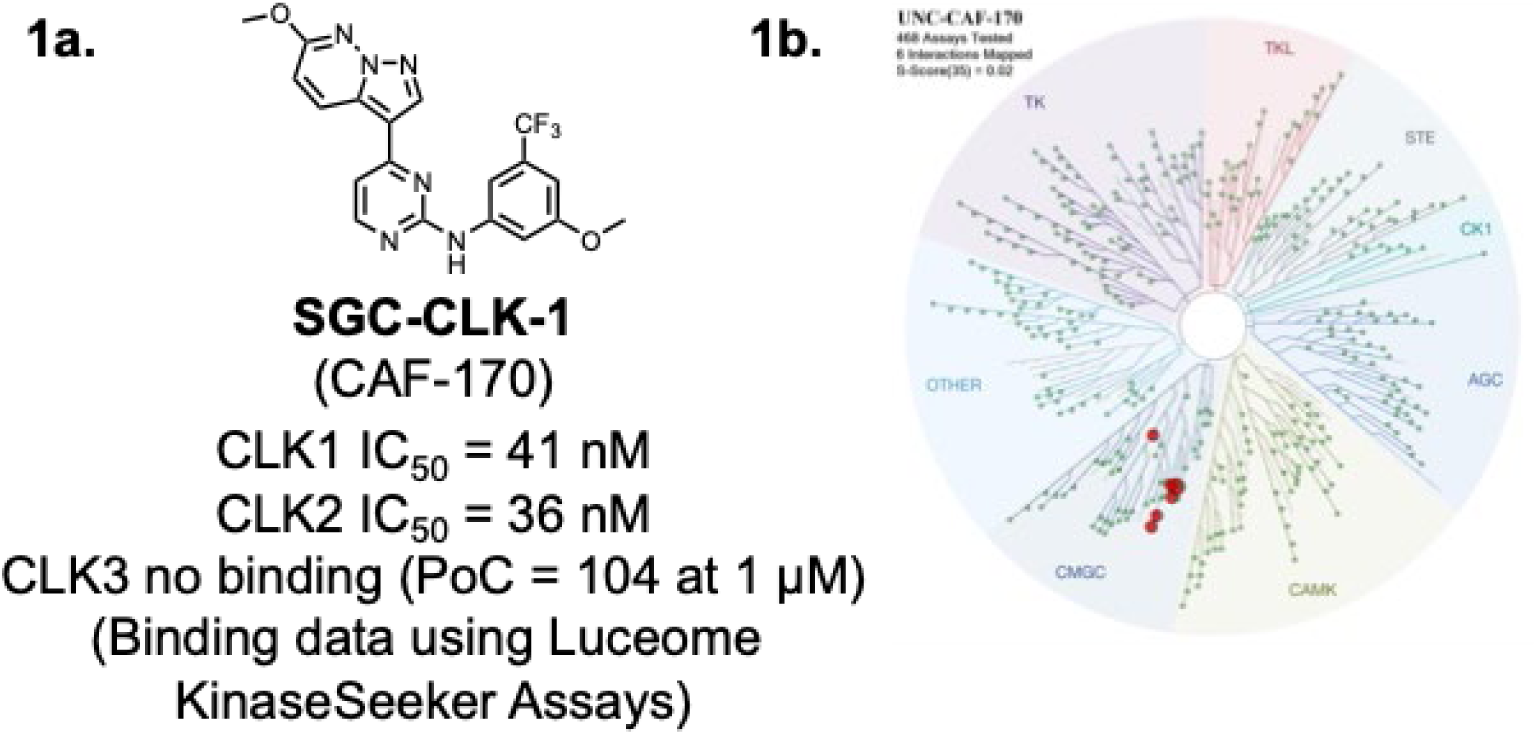
**SGC-CLK-1 (CAF-170): a**. SGC-CLK-1 structure **b**. SGC-CLK-1 KINOME*scan*® selectivity data visualized on the kinase phylogenetic tree.

The nine kinases that bound tightest to SGC-CLK-1 in the KINOMEscan® panel, along with family member CLK3 are listed in **Table 2**. To validate the binding data in an enzymatic assay, we generated enzyme IC_50_ inhibition data using commercially available assays (if available). SGC-CLK-1 is a potent inhibitor of CLK1, CLK2, and CLK4 (IC_50_ values, 13 nM, 4 nM, and 46 nM, respectively). CLK3 is inhibited more poorly, with IC_50_ = 363 nM. SGC-CLK-1 is also an inhibitor of HIPK1, HIPK2, and STK16. The NanoBRET in cell target engagement assay can be used to measure the binding to a kinase of interest in a cellular context. Thus, we employed that technology here to investigate cellular potency and selectivity. SGC-CLK-1 binds to CLK1, CLK2, and CLK4 with cellular IC_50_ values below 200 nM, with the CLK2 cellular affinity the best, at 58 nM (**Figure 2**). The compound does show some binding in cells to STK16, but only partial inhibition was achieved (Supplemental Table S1).

**Figure 2:**
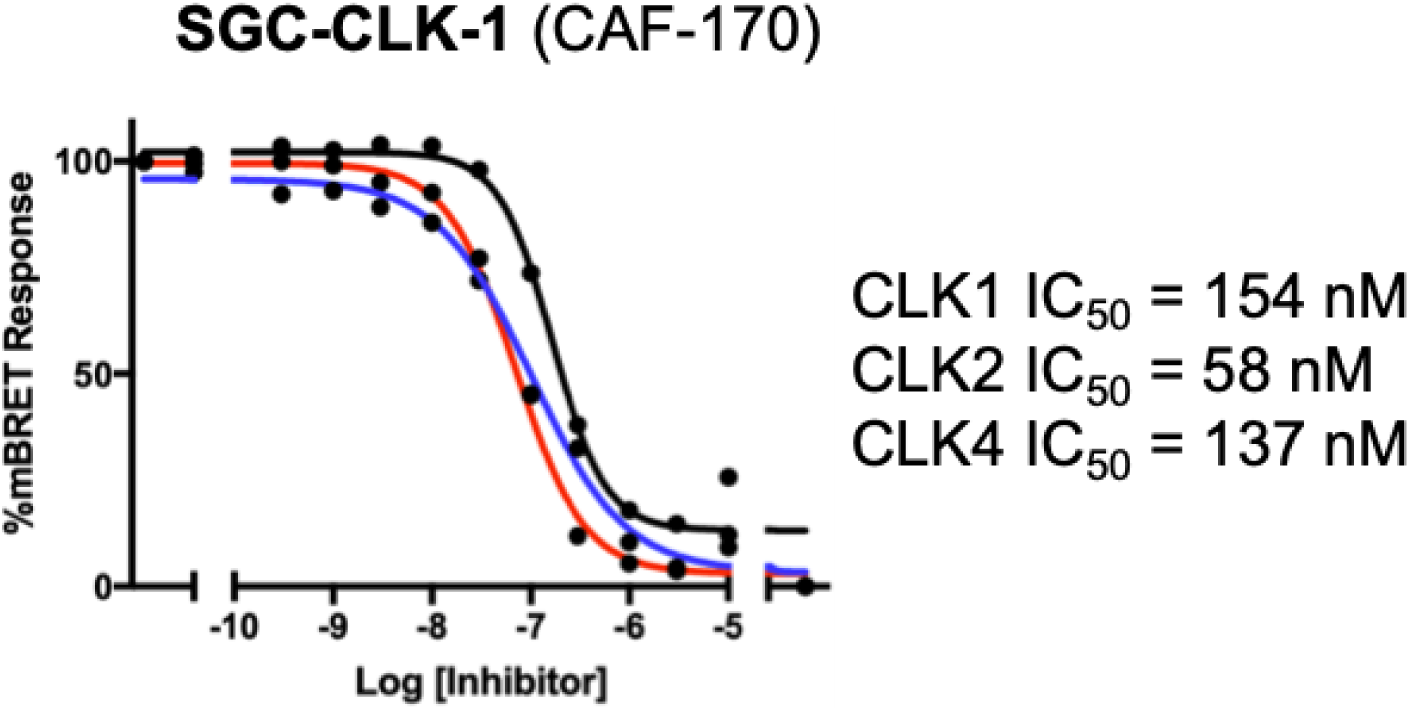
SGC-CLK-1 (CAF-170) binds to CLK1, CLK2, and CLK4 in cells.

**Table 2:** SGC-CLK-1 Binding, Enzymatic and Cellular Target Engagement characterization data.

| *Kinase* | % Control (1μM)<sup>a</sup> | IC<sub>50</sub> (nM)<sup>b</sup> | Cellular IC<sub>50</sub> (nM)<sup>c</sup> |
| --- | --- | --- | --- |
| CLK1 | 8 | 13 | 154 |
| CLK4 | 13 | 46 | 137 |
| CLK2 | 16 | 4 | 58 |
| ERK8 | 21 | nt | nt<sup>d</sup> |
| HIPK1 | 31 | 50 | >10,000 |
| HIPK2 | 34 | 42 | >10,000 |
| NEK7 | 40 | nt | nt<sup>d</sup> |
| PIP5K2B | 50 | nt<sup>e</sup> | nt<sup>d</sup> |
| STK16 | 53 | 49 | 167<sup>f</sup> |
| CLK3 | 92 | 363 | nt<sup>d</sup> |

a. % Control data was generated at Eurofins DiscoverX at a concentration of 1 μM
b. Enzymatic IC<sub>50</sub> values were generated at Eurofins
c. Cellular IC<sub>50</sub> values were generated in a NanoBRET assay
d. NanoBRET assay not available
e. No commercial enzyme assay available
f. NanoBRET showed only partial inhibition

In parallel, a structurally related compound, CAF-225 (renamed as SGC-CLK-1N), was designed and characterized as a negative control compound (**Figure 3a**). The key feature is the addition of a methyl group at the 6-position of the pyrimidine. In the KINOME*scan*® panel, CAF-225 did not bind to any kinases with PoC < 45. In the CLK Luceome KinaseSeeker™ CLK binding assays, CAF-225 showed 100% activity remaining for CLK1, CLK2, and CLK4 when screened at a concentration of 1 µM, indicating no binding to these targets at this concentration. CAF-225 was screened in enzymatic assays, and showed no inhibition of CLK1, CLK2, CLK3, CLK4 and a small panel of closely related targets, including the DYRK and HIPK families (Fig 3a, Supplemental Table S2). Finally, the negative control CAF-225 was inactive in the CLK1, CLK2, and CLK4 NanoBRET in-cell target engagement assays (**Figure 3b**).

**Figure 3:**
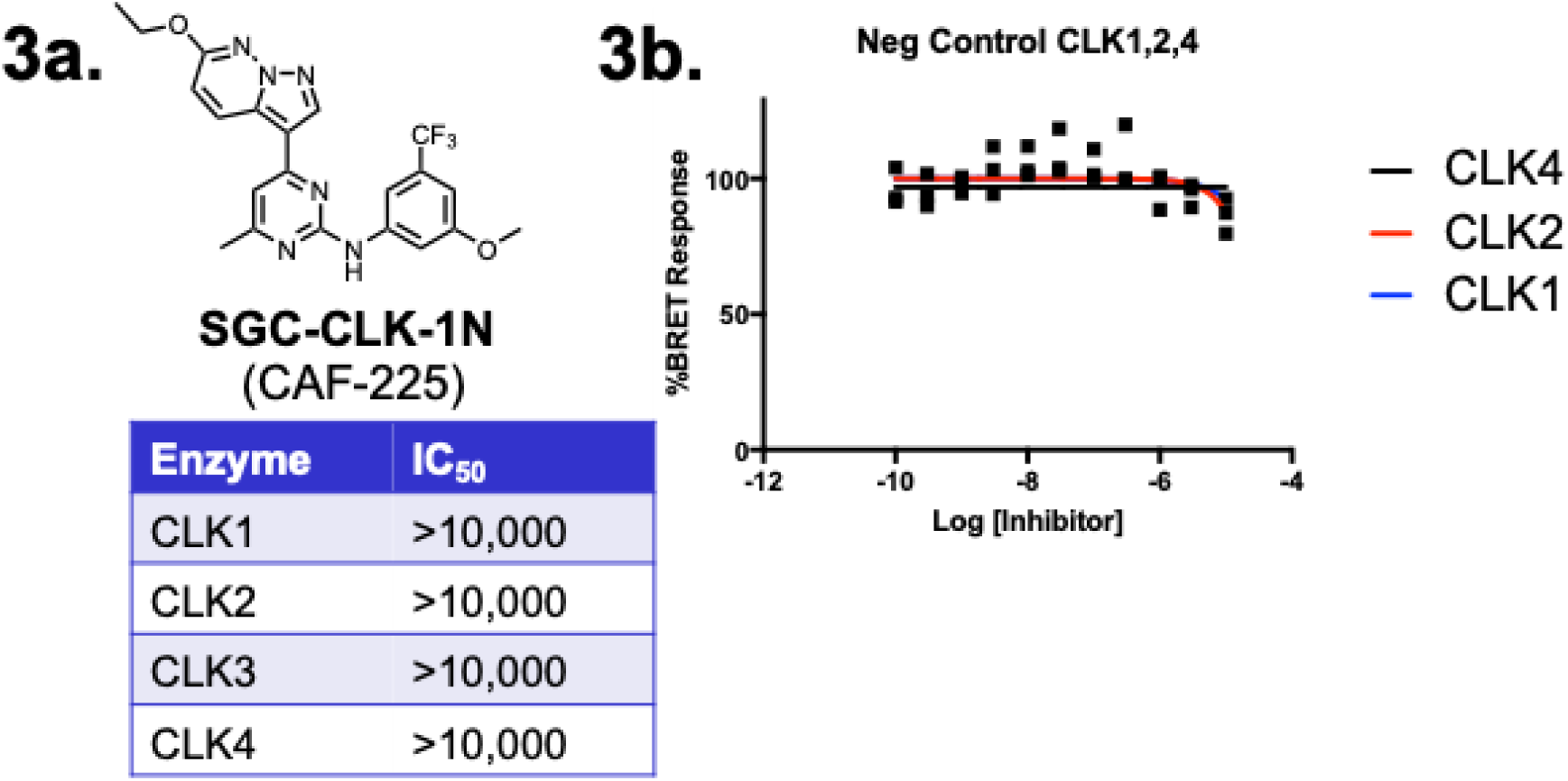
**3a**. **Structure of CLK negative control SGC-CLK-1N (CAF-225).** SGC-CLK-1N is inactive in CLK1, CLK2, CLK3, and CLK3 enzyme assays (Eurofins) **3b**. SGC-CLK-1N is inactive in the CLK1, CLK2, and CLK4 NanoBRET in cell target engagement assays.

### SGC-CLK-1 is a type 1 kinase inhibitor

Other CLK probes are available, but SGC-CLK-1 belongs to a structurally distinct chemical series. The availability of multiple chemotypes can be useful for elucidation of function as different scaffolds often have different patterns of off targets and different properties. In **Figure 4** we present the binding mode of SGC-CLK-1 alongside other CLK scaffolds.

**Figure 4:**
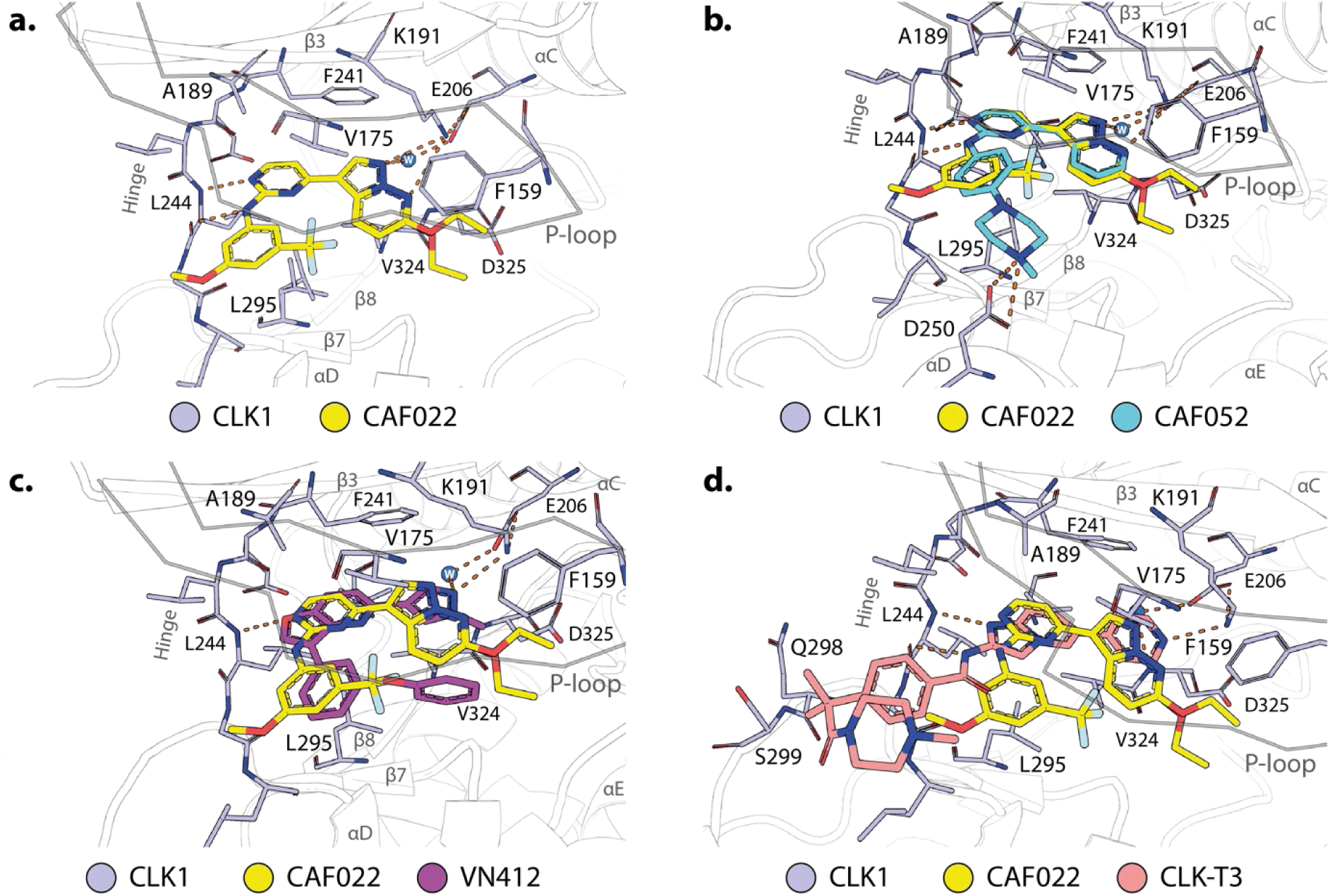
Structural analysis of CAF022 in comparison with other CLK inhibitors and chemical probes. **a.** Crystal structure of the CLK1 (gray)-CAF022 (yellow) complex (pdb 6ZLN). Highlighted are amino acids of CLK1 that directly interact with CAF022. Hydrogen bonds are indicated by orange dashes. For the ethyl ether of CAF022 two alternative conformations were modeled. **b-d**. Overlay of the binding modes of CAF022 (yellow) and **b**. CAF052 (cyan, pdb 7AK3), **c** VN412 (magenta, pdb 6I5H), or **d**. CLK-T3 (red, pdb 6RAA).

The crystal structure of the ethoxy analogue of CAF170, CAF022, has been recently published [45], highlighting the impact of the interaction of the CLK1 DFG-1 residue V324 with the pyrazolo[1,5-b]pyridazine (**Figure 4a**). The van-der-Waals interactions of this residue enable the correct positioning of the heterocycle to form hydrogen bonds with K191 and a structurally conserved water molecule. The importance of these back pocket interactions was confirmed by mutation studies of the V324 residue to alanine which resulted in a reduction the affinity by orders of magnitude [45]. Intriguingly, the ethoxy moiety showed two distinct conformation and was subsequently modeled with two alternative conformations. This suggests a lack of strong direct interaction of the ethoxy group but could potentially contribute to the strong selectivity of this compound series for CLK1/2/4. CAF022 forms further typical hydrogen bonds with the hinge via the backbone of L244. The 3-methoxy-5-(trifluoromethyl)phenyl engages in van-der-Waals interactions with the p-loop of CLK1 and thus contributing further to the potency of this compound. Interestingly, a chemically similar analog, CAF052, shares key interactions seen with CAF022 in complex with CLK1 (**Figure 4b**). This compound also shows high affinity towards CLK1 but lacks the selectivity for this protein family. For example, CAF052 has been recently identified as a potent ERK3 kinase binder [46] and the available KINOME*scan*® data of the trifluoromethyl derivative of CAF052, GW779439X [47], confirmed the promiscuous nature of these piperazine derivates. The solvent directed piperazine of CAF052 forms an electrostatic interaction with the carboxylic acid of D250. While the untypically strong back pocket interactions of CAF022 are a key driver of the potency for CLKs, the front pocket interactions of CAF052 are most likely not limited to CLK1 and therefore limit its selectivity. The strong back pocket interactions of CAF022 are also conserved in two other, highly selective CLK inhibitors, VN412 [30] and CLK-T3 [31].

Analogous to CAF022, both VN412 (**Figure 4c**) as well as CLK-T3 (**Figure 4d**) form direct van-der-Waals interaction with V324 and form similar hydrogen bonds with K191 and a conserved back pocket water molecule. Taken together, these structural models suggest that CAF022, analogous to CLK-T3 and VN412 exploits the non-conserved back pocket of CLK1 to gain selectivity over other kinases. This binding mode results in an affinity gain particular for the CLKs while avoiding extensive ATP mimetic kinase hinge contacts.

Thus, based on kinome wide screening, a panel of in vitro binding and enzymatic assays, NanoBRET in cell target engagement assays, and crystal structure and modeling data with the ethoxy analogue CAF-022, we show that CAF-170 is a unique CLK inhibitor suitable for exploring elements of CLK biology in cells.

### CAF-170 is active against multiple cancer types

As others have linked CLK activity to cancer cell growth, we screened CAF-170 (SGC-CLK-1) in the ProQinase CL100 cancer cell line panel. IC_50_ values for cell-proliferation were determined for 100 different cancer cell lines using a cell -titer glow assay format (full data set is on Supplemental table). SCG-CLK-1 has an IC_50_ value for inhibiting proliferation <250 nM for 27/100 cell lines tested. **Figure 5a** shows the IC_50_ values in subsets grouped by tissue type. As CLK2 showed the most potent inhibition with CAF-170, we focused on CLK2 expression to determine the best cancer type to target.

**Figure 5.**
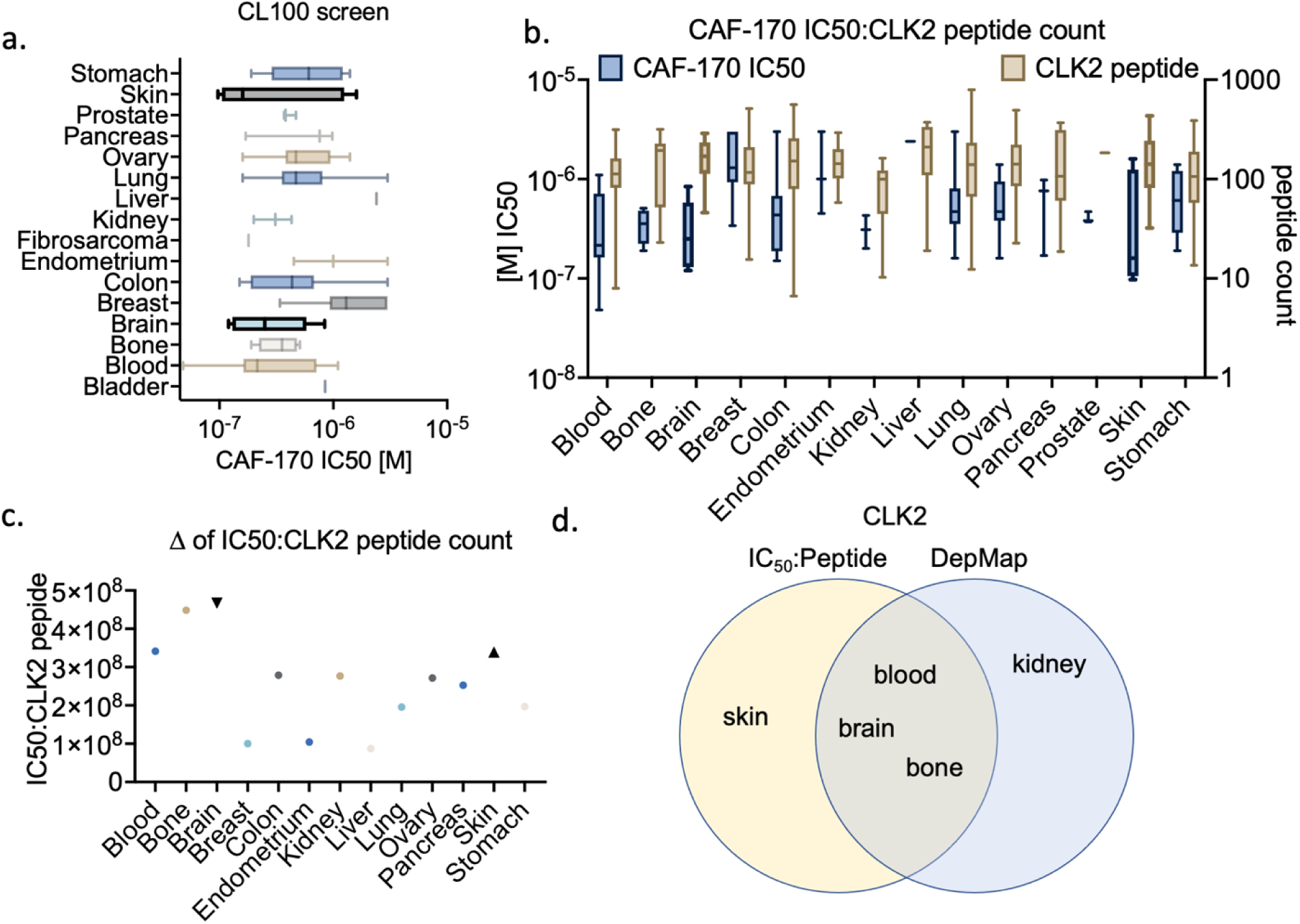
**a**. CL100 screen of denoted cancers by organ to determine IC_50_ of CAF-170 (SGC-CLK-1). **b**. CCLE quantification of CLK2 peptide abundance (tan boxplot) as compared to IC_50_ (blue boxplot from 4a) by cancer type. **c**. ratio of IC_50_ to CLK2 peptide abundance where a higher score suggests greater CLK2 dependence. d. DepMap and IC_50_:peptide ratio Venn diagram for overlapping CLK2 dependent cancers.

Using mass spectrometry data from the CCLE lines to determine the peptide abundance of CLK2 in each cancer by organ, as RNAseq data and protein abundance of splicing factors does not always correlate in a 1:1 fashion, we observed that some cancer types – blood, bone, brain, and skin – had an inverse relationship of low IC_50_ to high CLK2 peptide count (**Figure 5b**). To better define this relationship by cancer type, we calculated the average IC_50_ and CLK2 peptide abundance by cancer type and determined their IC_50_:peptide ratio. Confirming our observations in **Figure 5b**, we found blood, bone, brain, and skin with the highest ratios giving us potential cancers with a CLK2 dependency that could be targeted with a CLK2 inhibitor (**Figure 5c**). To test this idea, we went to The Cancer Dependency Map (DepMap) data to determine which cancers by organ have dependencies on single genes through both RNAi and CRIPSR screens. DepMap predicted CLK2 as a dependency for neuroblastoma (various), EWS_FLI (bone), glioma (brain) and glioblastoma (brain), and hematopoietic and lymphoid (blood), among others (**Figure 5d**). This substantial overlap between the IC_50_:peptide ratio and DepMap in cancer type was the greatest for CLK2 as compared to the other CLK family members (**Supplemental Figure 1**).

### CLK2 inhibition via CAF-170 (SGC-CLK-1) redistributes CLK2 and SR proteins

To test the efficacy of our CLK inhibitor in cell lines, we chose cell lines that showed either moderate (MDA-MB-435) or maximum (U118-MG) sensitivity to CAF-170 in our screen (**Figure 5a**) and had similar growth rates (**Supplemental Fig 2a**). Treatment with CAF-170 showed a dose-dependent decrease in the growth of both U118-MG and MDA-MB-435 cells, with U118-MG cells being more sensitive, as predicted, with little effect of the negative control compound CAF-225 (**Figure 6a**). As others have shown that CLK inhibition broadly decreases SR phosphorylation (pSR), we tested for but did not detect significant pSR changes at our highest treatment condition of 1 µM CAF-170. At 5 µM, however, we observed a decrease in SR phosphorylation 30 minutes post CAF-170 treatment (**Supplemental Figure 2b,c**). This, however, did not explain the decrease in cell growth we noted at a 10-fold lower concentration. We therefore looked at the localization of both pSRs and CLK2 post CAF-170 treatment. Surprisingly, we observed significant changes in the cellular distribution of both pSRs and CLK2 in cells treated with 500nM treatment of CAF-170 over the course of 60 minutes (**Figure 6b-d**). We next tested if the effect of our CLK2 probe could be reversed after CAF-170 was removed from the media. U118-MG cells were treated with 500nM CAF-170 for 15 minutes, and the media was replaced by conditioned U118-MG media where pSR and CLK2 localization was determined every 15 minutes for an hour. As shown in **Figure 6e** and quantified in **Figure 6f**, pSRs and CLK2 could redistribute to a normal punctate pattern post CAF-170 washout. Since our CLK2 probe showed a decrease in cell growth at a much lower concentration than other published CLK compounds, we wanted to test if other CLK inhibitors also affected pSR and CLK2 localization at nM-range concentrations. 500nM treatment with the active CLK probes – MU1210 and T3 – as well as their negative controls – MU140 and DMSO – did not show any effect on pSR or CLK2 localization. In this way, our CLK2 modulator seems to have a distinct mechanism of action at lower doses as compared to other published CLK inhibitors.

**Figure 6.**
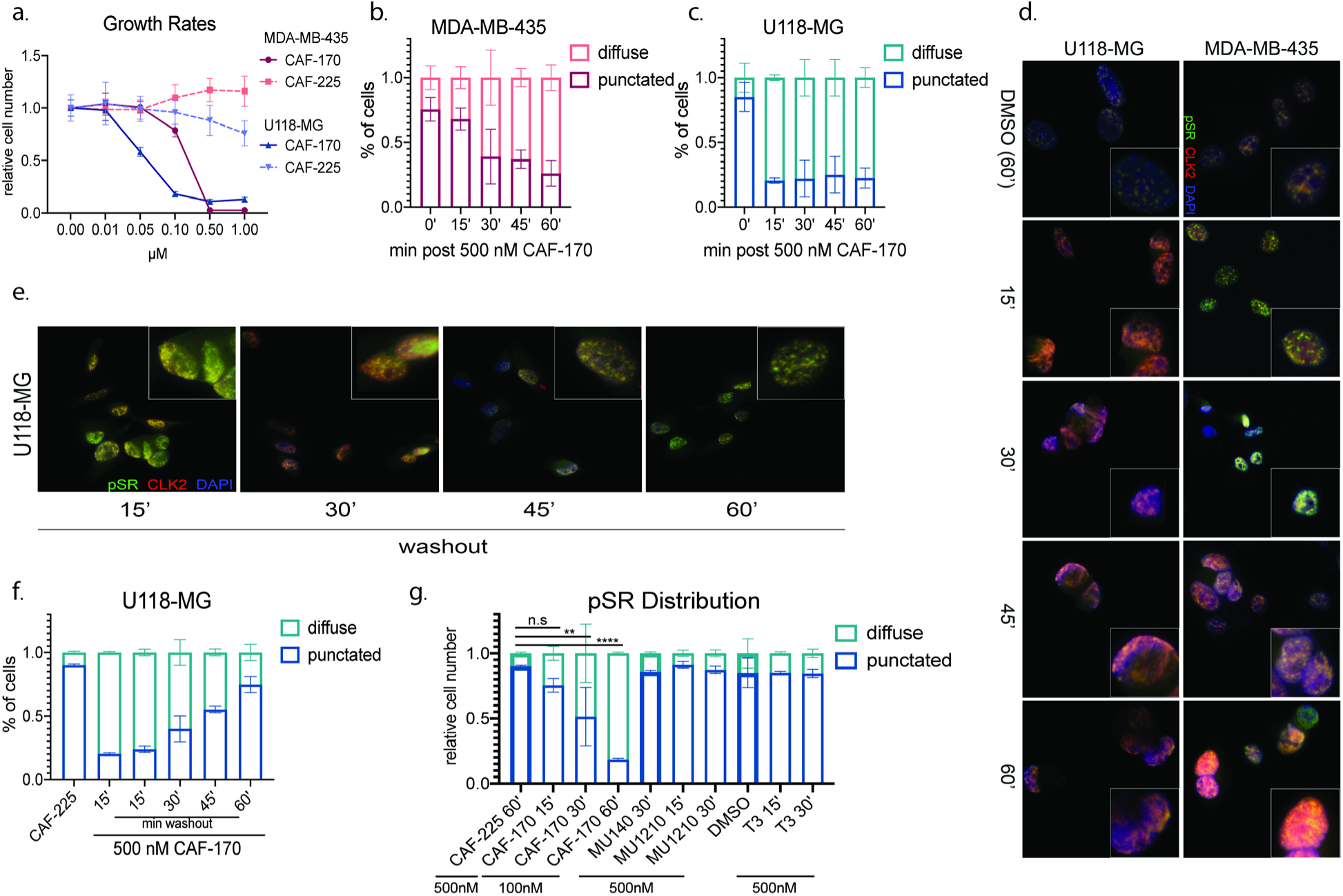
**a**. Crystal violet growth rates of U118-MG and MDA-MB-436 with denoted concentration of CAF-170 (SGC-CLK-1) or the negative control CAF-225 (SGC-CLK-1N) for 7 days. **b** and **c**. Immunofluorescent quantification of pSR localization at denoted timepoints and cell lines with 500 nM CAF-170 (SGC-CLK-1). **d** and **e**. Representative images of c. and f. quantification, respectively, in U118-MG cells. **f**. IF quantification of pSR localization post conditioned media washout in U118-MG cells. **g**. pSR localization via IF in denoted CLK inhibitors at either 500 or 100 nM concentrations for the times written. The bold bars depict the negative control for the respective CLK inhibitor.

## 4. Discussion

In this work, we show that SGC-CLK-1 (CAF-170) is a potent small molecule inhibitor of CLK1, 2 and 4 family members – with the greatest inhibition for CLK2. As the CLKs have garnered recent interest as potential therapeutic targets for osteoarthritis, solid tumors, neurodegenerative diseases, and drug refractory malignancies, a diverse set of tool compounds are necessary to better understand the role of these regulatory splicing kinases.

Using the *in vitro* CL100 cancer cell line panel IC_50_ screen, CCLE peptide abundance, and DepMap dependency information, we found that brain, skin, blood, and bone cancers show the lowest IC_50_ for CLK inhibition with the highest expression and greatest dependency for CLK2. We further confirm this in both a skin and brain cell line, where we find a dose-dependent decrease in growth with CAF-170 (SGC-CLK-1) as compared to our negative control compound CAF-225 (SGC-CLK-1N). While previous CLK inhibitors have focused on the phosphorylation status of serine/arginine rich (SR) proteins – a target of the CLK kinase family – we find that a high concentration of our probe (5 µM) decreases pSR abundance, but at a lower concentration of 500 nM this inhibitor disrupts pSR protein and CLK2 localization. The high dose effect on pSR phosphorylation is probably due to inhibition of all CLK isoforms (or other kinase targets) at the 5 µM concentration. The low dose phenotype, however, seems to be specific to our probe, as compared to other published probes, and warrants further exploration. Finally, we show that this compound can be washed out, which makes it a useful tool compound for live imaging experiments in the future to determine the localization (low dose) vs. kinase function (high dose) of the CLK family members.

As splicing modulators have become more prevalent in clinical use, a better understanding of the full role of these splicing factors and kinases are needed. CAF-170 (SGC-CLK-1) may be able to shed some light on the function of CLK family members with its non-canonical role of CLK and pSR mislocalization at low concentration vs its canonical role of phosphorylation modulation at higher doses. Future studies are needed to complete our understanding of this biologically important family and the roles they play in a diverse set of malignancies.

## Supporting information

Supporting Information for CLK probe

## Author contributions

DHD, DMT, CIW, and WZ conceived the project. DMT, CIW, MS, CAF, CdeS, FK, PS, AP, and MAH performed the experiments. MS, SK, RBR and SYC provided general advice. Manuscript writing – Original Draft, DMT and DHD, Writing – Review & Editing, MS, XS, CAF, AG, RI, ML, BH, MAH, WZ, RZ, SK, RBR, and SYC. Funding Acquisition, DHD, RZ, SYC, and DMT.

## Acknowledgements

Funding for this project was provided by these grants from the National Institutes of Health: 1R44TR001916-02 to Luceome Biotechnologies and the SGC-UNC, NS125318 to Northwestern University (Shi-Yuan Cheng_),_ and NIH K00 CA234799 to Northwestern University (Deanna Tiek). We are grateful for support from the Structural Genomics Consortium (SGC), a registered charity (no. 1097737) that receives funds from Bayer AG, Boehringer Ingelheim, Bristol Myers Squibb, Genentech, Genome Canada through Ontario Genomics Institute [OGI196], EU/EFPIA/OICR/McGill/KTH/Diamond Innovative Medicines Initiative 2 Joint Undertaking [EUbOPEN grant 875510], Janssen, Merck KGaA (aka EMD in Canada and USA), Pfizer, and Takeda. We thank Brandie Ehrmann and Diane Weatherspoon of the University of North Carolina’s Department of Chemistry Mass Spectrometry Core Laboratory for assistance with mass spectrometry analysis.

## Reference

1. Roskoski, R., Jr. Properties of FDA-approved small molecule protein kinase inhibitors: A 2022 update. Pharmacol Res 2022, 175, 106037, doi:10.1016/j.phrs.2021.106037.

2. Attwood, M.M.; Fabbro, D.; Sokolov, A.V.; Knapp, S.; Schioth, H.B. Trends in kinase drug discovery: targets, indications and inhibitor design. Nat Rev Drug Discov 2021, 20, 839–861, doi:10.1038/s41573-021-00252-y.

3. Gallego-Paez, L.M.; Bordone, M.C.; Leote, A.C.; Saraiva-Agostinho, N.; Ascensao-Ferreira, M.; Barbosa-Morais, N.L. Alternative splicing: the pledge, the turn, and the prestige : The key role of alternative splicing in human biological systems. Hum Genet 2017, 136, 1015–1042, doi:10.1007/s00439-017-1790-y.

4. Lee, Y.; Rio, D.C. Mechanisms and Regulation of Alternative Pre-mRNA Splicing. Annu Rev Biochem 2015, 84, 291–323, doi:10.1146/annurev-biochem-060614-034316.

5. Bates, D.O.; Morris, J.C.; Oltean, S.; Donaldson, L.F. Pharmacology of Modulators of Alternative Splicing. Pharmacol Rev 2017, 69, 63–79, doi:10.1124/pr.115.011239.

6. Le, K.Q.; Prabhakar, B.S.; Hong, W.J.; Li, L.C. Alternative splicing as a biomarker and potential target for drug discovery. Acta Pharmacol Sin 2015, 36, 1212–1218, doi:10.1038/aps.2015.43.

7. Martin Moyano, P.; Nemec, V.; Paruch, K. Cdc-Like Kinases (CLKs): Biology, Chemical Probes, and Therapeutic Potential. Int J Mol Sci 2020, 21, doi:10.3390/ijms21207549.

8. Ren, P.; Lu, L.; Cai, S.; Chen, J.; Lin, W.; Han, F. Alternative Splicing: A New Cause and Potential Therapeutic Target in Autoimmune Disease. Front Immunol 2021, 12, 713540, doi:10.3389/fimmu.2021.713540.

9. McClorey, G.; Fletcher, S.; Wilton, S. Splicing intervention for Duchenne muscular dystrophy. Curr Opin Pharmacol 2005, 5, 529–534, doi:10.1016/j.coph.2005.06.001.

10. Pistoni, M.; Ghigna, C.; Gabellini, D. Alternative splicing and muscular dystrophy. RNA Biol 2010, 7, 441–452, doi:10.4161/rna.7.4.12258.

11. Li, D.; McIntosh, C.S.; Mastaglia, F.L.; Wilton, S.D.; Aung-Htut, M.T. Neurodegenerative diseases: a hotbed for splicing defects and the potential therapies. Transl Neurodegener 2021, 10, 16, doi:10.1186/s40035-021-00240-7.

12. Bonnal, S.C.; Lopez-Oreja, I.; Valcarcel, J. Roles and mechanisms of alternative splicing in cancer - implications for care. Nat Rev Clin Oncol 2020, 17, 457–474, doi:10.1038/s41571-020-0350-x.

13. Di, C.; Syafrizayanti; Zhang, Q.; Chen, Y.; Wang, Y.; Zhang, X.; Liu, Y.; Sun, C.; Zhang, H.; Hoheisel, J.D. Function, clinical application, and strategies of Pre-mRNA splicing in cancer. Cell Death Differ 2019, 26, 1181–1194, doi:10.1038/s41418-018-0231-3.

14. Lindberg, M.F.; Meijer, L. Dual-Specificity, Tyrosine Phosphorylation-Regulated Kinases (DYRKs) and cdc2-Like Kinases (CLKs) in Human Disease, an Overview. Int J Mol Sci 2021, 22, doi:10.3390/ijms22116047.

15. Schneider-Poetsch, T.; Chhipi-Shrestha, J.K.; Yoshida, M. Splicing modulators: on the way from nature to clinic. J Antibiot (Tokyo) 2021, 74, 603–616, doi:10.1038/s41429-021-00450-1.

16. Melnyk, J.E.; Steri, V.; Nguyen, H.G.; Hann, B.; Feng, F.Y.; Shokat, K.M. The splicing modulator sulfonamide indisulam reduces AR-V7 in prostate cancer cells. Bioorganic & medicinal chemistry 2020, 28, 115712, doi:10.1016/j.bmc.2020.115712.

17. Bowler, E.; Porazinski, S.; Uzor, S.; Thibault, P.; Durand, M.; Lapointe, E.; Rouschop, K.M.A.; Hancock, J.; Wilson, I.; Ladomery, M. Hypoxia leads to significant changes in alternative splicing and elevated expression of CLK splice factor kinases in PC3 prostate cancer cells. BMC Cancer 2018, 18, 355, doi:10.1186/s12885-018-4227-7.

18. Braun, C.J.; Stanciu, M.; Boutz, P.L.; Patterson, J.C.; Calligaris, D.; Higuchi, F.; Neupane, R.; Fenoglio, S.; Cahill, D.P.; Wakimoto, H.; Agar, N.Y.R.; Yaffe, M.B.; Sharp, P.A.; Hemann, M.T.; Lees, J.A. Coordinated Splicing of Regulatory Detained Introns within Oncogenic Transcripts Creates an Exploitable Vulnerability in Malignant Glioma. Cancer Cell 2017, 32, 411–426 e411, doi:10.1016/j.ccell.2017.08.018.

19. Tiek, D.M.; Khatib, S.A.; Trepicchio, C.J.; Heckler, M.M.; Divekar, S.D.; Sarkaria, J.N.; Glasgow, E.; Riggins, R.B. Estrogen-related receptor beta activation and isoform shifting by cdc2-like kinase inhibition restricts migration and intracranial tumor growth in glioblastoma. FASEB J 2019, 33, 13476–13491, doi:10.1096/fj.201901075R.

20. Tiek, D.M.; Erdogdu, B.; Razaghi, R.; Jin, L.; Sadowski, N.; Alamillo-Ferrer, C.; Hogg, J.R.; Haddad, B.R.; Drewry, D.H.; Wells, C.I.; Pickett, J.E.; Song, X.; Goenka, A.; Hu, B.; Goldlust, S.A.; Zuercher, W.J.; Pertea, M.; Timp, W.; Cheng, S.Y.; Riggins, R.B. Temozolomide-induced guanine mutations create exploitable vulnerabilities of guanine-rich DNA and RNA regions in drug-resistant gliomas. Sci Adv 2022, 8, eabn3471, doi:10.1126/sciadv.abn3471.

21. Oltean, S.; Bates, D.O. Hallmarks of alternative splicing in cancer. Oncogene 2014, 33, 5311–5318, doi:10.1038/onc.2013.533.

22. Hanahan, D.; Weinberg, R.A. Hallmarks of cancer: the next generation. Cell 2011, 144, 646–674, doi:10.1016/j.cell.2011.02.013.

23. Hanahan, D.; Weinberg, R.A. The hallmarks of cancer. Cell 2000, 100, 57–70, doi:10.1016/s0092-8674(00)81683-9.

24. Ladomery, M. Aberrant alternative splicing is another hallmark of cancer. Int J Cell Biol 2013, *2013*, 463-786, doi:10.1155/2013/463786.

25. Prasad, J.; Manley, J.L. Regulation and substrate specificity of the SR protein kinase Clk/Sty. Mol Cell Biol 2003, 23, 4139–4149, doi:10.1128/MCB.23.12.4139-4149.2003.

26. Ngo, J.C.; Chakrabarti, S.; Ding, J.H.; Velazquez-Dones, A.; Nolen, B.; Aubol, B.E.; Adams, J.A.; Fu, X.D.; Ghosh, G. Interplay between SRPK and Clk/Sty kinases in phosphorylation of the splicing factor ASF/SF2 is regulated by a docking motif in ASF/SF2. Mol Cell 2005, 20, 77–89, doi:10.1016/j.molcel.2005.08.025.

27. Mueller, W.F.; Hertel, K.J. The role of SR and SR-related proteins in pre-mRNA splicing. In RNA Binding Proteins 2012, 27–46.

28. Bullock, A.N.; Das, S.; Debreczeni, J.E.; Rellos, P.; Fedorov, O.; Niesen, F.H.; Guo, K.; Papagrigoriou, E.; Amos, A.L.; Cho, S.; Turk, B.E.; Ghosh, G.; Knapp, S. Kinase domain insertions define distinct roles of CLK kinases in SR protein phosphorylation. Structure 2009, 17, 352–362, doi:10.1016/j.str.2008.12.023.

29. Qin, Z.; Qin, L.; Feng, X.; Li, Z.; Bian, J. Development of Cdc2-like Kinase 2 Inhibitors: Achievements and Future Directions. Journal of medicinal chemistry 2021, 64, 13191–13211, doi:10.1021/acs.jmedchem.1c00985.

30. Nemec, V.; Hylsova, M.; Maier, L.; Flegel, J.; Sievers, S.; Ziegler, S.; Schroder, M.; Berger, B.T.; Chaikuad, A.; Valcikova, B.; Uldrijan, S.; Drapela, S.; Soucek, K.; Waldmann, H.; Knapp, S.; Paruch, K. Furo[3,2-b]pyridine: A Privileged Scaffold for Highly Selective Kinase Inhibitors and Effective Modulators of the Hedgehog Pathway. Angew Chem Int Ed Engl 2019, 58, 1062–1066, doi:10.1002/anie.201810312.

31. Funnell, T.; Tasaki, S.; Oloumi, A.; Araki, S.; Kong, E.; Yap, D.; Nakayama, Y.; Hughes, C.S.; Cheng, S.G.; Tozaki, H.; Iwatani, M.; Sasaki, S.; Ohashi, T.; Miyazaki, T.; Morishita, N.; Morishita, D.; Ogasawara-Shimizu, M.; Ohori, M.; Nakao, S.; Karashima, M.; Sano, M.; Murai, A.; Nomura, T.; Uchiyama, N.; Kawamoto, T.; Hara, R.; Nakanishi, O.; Shumansky, K.; Rosner, J.; Wan, A.; McKinney, S.; Morin, G.B.; Nakanishi, A.; Shah, S.; Toyoshiba, H.; Aparicio, S. CLK-dependent exon recognition and conjoined gene formation revealed with a novel small molecule inhibitor. Nat Commun 2017, 8, 7, doi:10.1038/s41467-016-0008-7.

32. Sun, Q.Z.; Lin, G.F.; Li, L.L.; Jin, X.T.; Huang, L.Y.; Zhang, G.; Yang, W.; Chen, K.; Xiang, R.; Chen, C.; Wei, Y.Q.; Lu, G.W.; Yang, S.Y. Discovery of Potent and Selective Inhibitors of Cdc2-Like Kinase 1 (CLK1) as a New Class of Autophagy Inducers. Journal of medicinal chemistry 2017, 60, 6337–6352, doi:10.1021/acs.jmedchem.7b00665.

33. Tam, B.Y.; Chiu, K.; Chung, H.; Bossard, C.; Nguyen, J.D.; Creger, E.; Eastman, B.W.; Mak, C.C.; Ibanez, M.; Ghias, A.; Cahiwat, J.; Do, L.; Cho, S.; Nguyen, J.; Deshmukh, V.; Stewart, J.; Chen, C.W.; Barroga, C.; Dellamary, L.; Kc, S.K.; Phalen, T.J.; Hood, J.; Cha, S.; Yazici, Y. The CLK inhibitor SM08502 induces anti-tumor activity and reduces Wnt pathway gene expression in gastrointestinal cancer models. Cancer Lett 2020, 473, 186–197, doi:10.1016/j.canlet.2019.09.009.

34. Bossard, C.; Cruz, N.; Chiu, K.; Eastman, B.; Mak, C.C.; Kc, S.; Bucci, G.; Stewart, J.; Phalen, T.J.; Cha, S. Abstract 5691: SM08502, a novel, small-molecule CDC-like kinase (CLK) inhibitor, demonstrates strong antitumor effects and Wnt pathway inhibition in castration-resistant prostate cancer (CRPC) models. Cancer Research 2020, *80*, 5691–5691, doi:10.1158/1538-7445.Am2020-5691.

35. Hood, J.; Wallace David, M.; Kc Sunil, K.; Yazici, Y.; Swearingen, C.; Dellamary Luis, A. 2-(1H-INDAZOL-3-YL)-3H-IMIDAZO[4,5-C]PYRIDINES AND THEIR ANTI-INFLAMMATORY USES THEREOF. WO 2017/079759 A1, 2016/11/07 2017.

36. Kc Sunil, K. PROCESS FOR PREPARING N-(5-(3-(7-(3-FLUOROPHENYL)-3H-IMIDAZO[4,5-C]PYRIDIN-2-YL)-1H-INDAZOL-5-YL)PYRIDIN-3-YL)-3-METHYLBUTANAMIDE. WO 2017/210407 A1, 2017/06/01 2017.

37. Ogawa, S.; Miyake, H.; Makishima, H.; Nannya, Y.; Ochi, Y.; Satoh, Y.; Mizutani, A.; Morishita, D.; Yoda, A. CTX-712, a Novel Clk Inhibitor Targeting Myeloid Neoplasms with SRSF2 Mutation. Blood 2019, 134, 404–404, doi:10.1182/blood-2019-131559.

38. Kim, H.; Choi, K.; Kang, H.; Lee, S.Y.; Chi, S.W.; Lee, M.S.; Song, J.; Im, D.; Choi, Y.; Cho, S. Identification of a novel function of CX-4945 as a splicing regulator. Plos One 2014, 9, e94978, doi:10.1371/journal.pone.0094978.

39. Lee, J.Y.; Yun, J.S.; Kim, W.K.; Chun, H.S.; Jin, H.; Cho, S.; Chang, J.H. Structural Basis for the Selective Inhibition of Cdc2-Like Kinases by CX-4945. Biomed Res Int 2019, 2019, 6125068, doi:10.1155/2019/6125068.

40. Riggs, J.R.; Nagy, M.; Elsner, J.; Erdman, P.; Cashion, D.; Robinson, D.; Harris, R.; Huang, D.; Tehrani, L.; Deyanat-Yazdi, G.; Narla, R.K.; Peng, X.; Tran, T.; Barnes, L.; Miller, T.; Katz, J.; Tang, Y.; Chen, M.; Moghaddam, M.F.; Bahmanyar, S.; Pagarigan, B.; Delker, S.; LeBrun, L.; Chamberlain, P.P.; Calabrese, A.; Canan, S.S.; Leftheris, K.; Zhu, D.; Boylan, J.F. The Discovery of a Dual TTK Protein Kinase/CDC2-Like Kinase (CLK2) Inhibitor for the Treatment of Triple Negative Breast Cancer Initiated from a Phenotypic Screen. Journal of medicinal chemistry 2017, 60, 8989–9002, doi:10.1021/acs.jmedchem.7b01223.

41. Shi, Y.; Park, J.; Lagisetti, C.; Zhou, W.; Sambucetti, L.C.; Webb, T.R. A triple exon-skipping luciferase reporter assay identifies a new CLK inhibitor pharmacophore. Bioorg Med Chem Lett 2017, 27, 406–412, doi:10.1016/j.bmcl.2016.12.056.

42. Feoktistova, M.; Geserick, P.; Leverkus, M. Crystal Violet Assay for Determining Viability of Cultured Cells. Cold Spring Harb Protoc 2016, 2016, pdb prot087379, doi:10.1101/pdb.prot087379.

43. Jester, B.W.; Cox, K.J.; Gaj, A.; Shomin, C.D.; Porter, J.R.; Ghosh, I. A coiled-coil enabled split-luciferase three-hybrid system: applied toward profiling inhibitors of protein kinases. J Am Chem Soc 2010, 132, 11727–11735, doi:10.1021/ja104491h.

44. Jester, B.W.; Gaj, A.; Shomin, C.D.; Cox, K.J.; Ghosh, I. Testing the promiscuity of commercial kinase inhibitors against the AGC kinase group using a split-luciferase screen. Journal of medicinal chemistry 2012, 55, 1526–1537, doi:10.1021/jm201265f.

45. Schroder, M.; Bullock, A.N.; Fedorov, O.; Bracher, F.; Chaikuad, A.; Knapp, S. DFG-1 Residue Controls Inhibitor Binding Mode and Affinity, Providing a Basis for Rational Design of Kinase Inhibitor Selectivity. Journal of medicinal chemistry 2020, 63, 10224–10234, doi:10.1021/acs.jmedchem.0c00898.

46. Schroder, M.; Filippakopoulos, P.; Schwalm, M.P.; Ferrer, C.A.; Drewry, D.H.; Knapp, S.; Chaikuad, A. Crystal Structure and Inhibitor Identifications Reveal Targeting Opportunity for the Atypical MAPK Kinase ERK3. Int J Mol Sci 2020, 21, doi:10.3390/ijms21217953.

47. Drewry, D.H.; Wells, C.I.; Andrews, D.M.; Angell, R.; Al-Ali, H.; Axtman, A.D.; Capuzzi, S.J.; Elkins, J.M.; Ettmayer, P.; Frederiksen, M.; Gileadi, O.; Gray, N.; Hooper, A.; Knapp, S.; Laufer, S.; Luecking, U.; Michaelides, M.; Muller, S.; Muratov, E.; Denny, R.A.; Saikatendu, K.S.; Treiber, D.K.; Zuercher, W.J.; Willson, T.M. Progress towards a public chemogenomic set for protein kinases and a call for contributions. Plos One 2017, 12, e0181585, doi:10.1371/journal.pone.0181585.

