## Supporting Information for CLK probe for "SGC-CLK-1 (CAF-170) a chemical probe for the Cdc2-Like kinases CLK1, CLK2, and CLK4"

**Table S1: CAF-170 Activity Profile**

| Kinase | % Inhibition  from Control (1µM) CAF-170 | Enzyme  IC_50_ (nM) | NanoBRET  (nM) |
| --- | --- | --- | --- |
| CLK1 | 8 | 13 | 165 |
| **CLK2** | **16** | **4** | **70** |
| CLK3 | 92 | 363 | nt** |
| CLK4 | 13 | 46 | 100 |
| HIPK1 | 31 | 50 | >10,000 |
| HIPK2 | 34 | 42 | >10,000 |
| NEK7 | 40 | nt | nt** |
| PIP5K2B | 50 | nt | nt** |
| STK16 | 53 | 49 | 167 |

**Table S2: CAF-225 Enzymatic Profile**

**Eurofins Enzymatic Screening**

| Kinase | CAF-225 IC_50_ (nM) |
| --- | --- |
| CLK1 | >10,000 |
| CLK2 | >10,000 |
| CLK3 | >10,000 |
| CLK4 | >10,000 |
| DYRK1A | >10,000 |
| DYRK1B | >10,000 |
| DYRK2 | >10,000 |
| DYRK3 | >10,000 |
| HIPK1 | >10,000 |
| HIPK2 | >10,000 |
| HIPK3 | >10,000 |
| HIPK4 | >10,000 |
| Pim-2 | >10,000 |

**Supplemental Figure 1: Related to Fig 5**

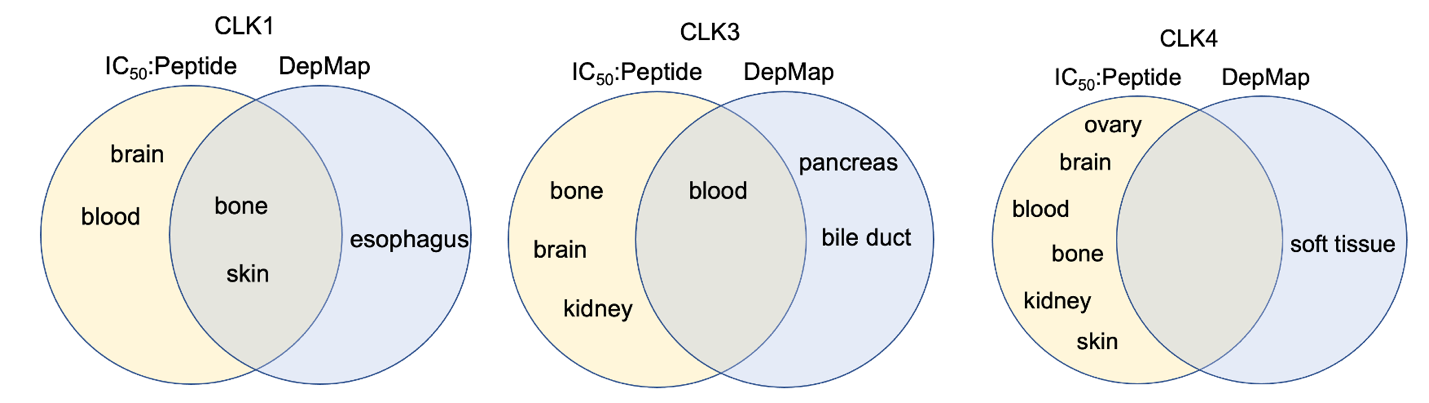

**Supplementary Figure 1. Venn Diagrams of CLK1,3 and 4 overlap between DepMap and CL100 screen.** IC50:Peptide and DepMap Venn Diagrams for the other 3 CLK members, where CLK2 has the highest overlap of predicted dependencies.

**Supplemental Figure 2. Related to Fig 6**

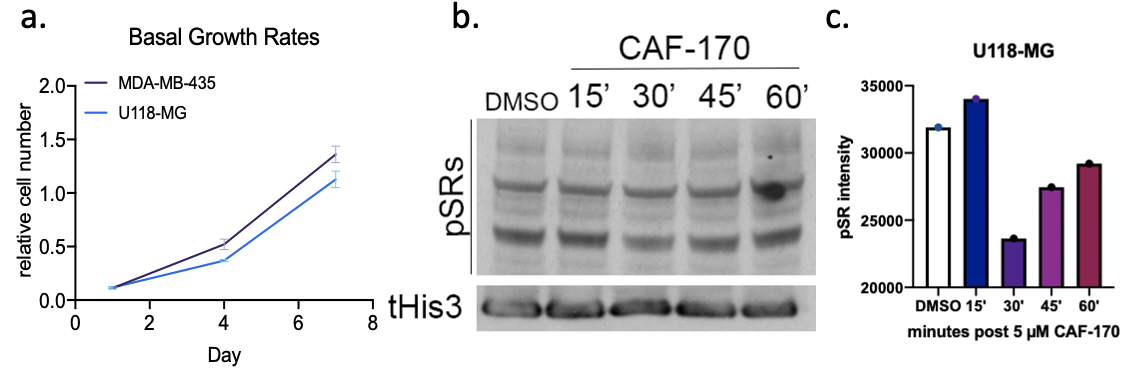

**Supplementary Figure 2. Changes in phosphorylation with high dose CAF-170.** A. Crystal violet assay to determine basal growth rates of denoted cells at denoted time points. B. Western blot of pSR proteins (1H4) in U118-MG cells post 5µM CAF-170 at denoted times (left), with c. quantification on the right.

**CLK Compounds synthesis procedures**

General Chemistry Information

All reagents and solvents, unless specifically stated, were used as obtained from their commercial sources without further purification. Air and moisture sensitive reactions were performed under an inert atmosphere using nitrogen in a previously oven-dried or flame-dried reaction flask, and addition of reagents were done using a syringe. All microwave (MW) reactions were carried out in a Biotage Initiator EXP US 400W microwave synthesizer. Thin layer chromatography (TLC) analyses were performed using 200 μm pre-coated sorbtech fluorescent TLC plates and spots were visualized using UV light. High resolution mass spectrometry samples were analyzed with a ThermoFisher Q Exactive HF-X (ThermoFisher, Bremen, Germany) mass spectrometer coupled with a Waters Acquity H-class liquid chromatograph system. Column chromatography was undertaken with a Biotage Isolera One instrument. Nuclear magnetic resonance (NMR) spectrometry was run on a varian Inova 400 MHz or Bruker Avance III 700 MHz spectrometer equipped with a TCI H-C/N-D 5 mm cryoprobe and data was processed using the MestReNova processor. Chemical shifts are reported in ppm with residual solvent peaks referenced as internal standard.

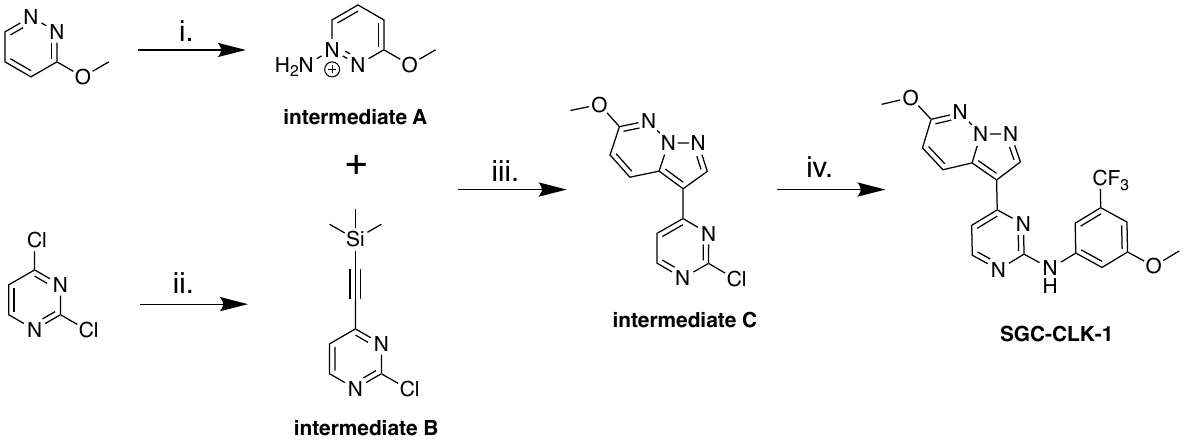

**Reagent and conditions:** i) Amino hydrogen sulphate, KHCO_3_, H_2_O, 80 C, 14 h; ii) KHCO_3_, KOH, H_2_O, DCM, r.t, 18 h; iii) ETMS, Pd(dppf)Cl_2_ · CH_2_Cl_2_, CuI, PPh_3_, TEA, THF, 70 C, 15 min; iv) TFA, tert-BuOH, 85 C, 15 h.

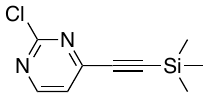

**2-chloro-4-((trimethylsilyl)ethynyl)pyrimidine (Intermediate B):**

2,4-dichloropyrimidine (1.00 g, 6.7 mmol, 1 eq.) and tetrahydrofuran (30 mL) degassed with nitrogen for 10-12 min, then ethynyltrimethylsilane (0.73 g, 7.4 mmol, 1.1 eq.) and triethylamine (0.74 g, 7.5 mmol, 1.1 eq.) added, degassed for another 5min, then the rest of reagents PdCl2(dppf)-CH2Cl2adduct (0.27 g, 0.34 mmol, 0.05 eq.), copper(I) iodide (0.13 g, 0.67 mmol, 0.10 eq.), triphenylphosphine (0.18 g, 0.67 mmol, 0.10 eq.) were added. Reaction was then refluxed for 15 min. Bulky solid formed. By TLC, all starting material was consumed (Hexanes:EtOAc, 85:15). Hexanes added to precipitated OPPh_3_, filtered and rinsed with EtOAc. The crude material was concentrated in the rotavap, thick brown liquid formed. Desired product obtained using flash chromatography using EtOAc in hexanes (0% to 20%). Concentrated in rotavapor, dried under vacuum overnight and desired product yielded as brown solid 770 mg (yield 50%), purity>90% by NMR.

^1^H NMR (400 MHz, DMSO-*d*_6_) δ ppm 0.27 (s, 9 H), 7.66 (d, *J*=5.1 Hz, 1 H), 8.80 (d, *J*=5.1 Hz, 1 H).

^13^C NMR (101 MHz, DMSO-*d*_6_) δ ppm 100.4, 102.1, 122.6, 151.4, 160.1, 161.2.

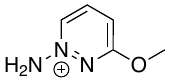

**1-amino-3-methoxypyridazin-1-ium (Intermediate A):**

Amino hydrogen sulphate (0.83 g, 7.39 mmol, 3.7 eq.) was dissolved in water (1.6 mL), KHCO_3_ (0.74 g, 7.39 mmol, 3.7 eq.) in water (1.0 mL), pH 5. Then 3-methoxypyridazine (0.22 g, 2.00 mmol, 1.00 eq.) was added in portion. Reaction was then stirred at 80 °C overnight. This crude is used as is for the next reaction without purification.

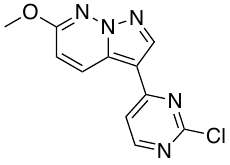

**3-(2-chloropyrimidin-4-yl)-6-methoxypyrazolo[1,5-b]pyridazine (Intermediate C):**

1-amino-3-methoxypyridazin-1-ium (0.20 g, 1.6 mmol, 1.5 eq.) from the previous step (pH = 1) was treated with saturated KHCO_3_ to bring the pH to 7. 2-chloro-4-((trimethylsilyl)ethynyl)pyrimidine (0.22 g, 1.04 mmol, 1.00 eq.) was dissolved in 1 mL of DCM (1 M), added in one portion over the crude. KOH (0. 320 g, 6.26 mmol, 6 eq.) was dissolved in H_2_O (5 mL) 1.0 M and was added in one portion over previous mixture. The reaction mixture was transformed dark read in color after 5-10 min. Reaction mixture was strongly stirred at r.t. for 22 h. The crude mixture was then quenched with water, extracted with DCM, and combined organic layers dried over anhydrous Na_2_SO_4_. Crude was dry loaded on a 10 g Biotage Sfar 60 um silica cartridge (Hexanes/EtOAc 70:30) and whitish/light pink solid yielded (0.095 g, 35% yield). LCMS [M+1] = 262, purity > 95%.

^1^H NMR (400 MHz, DMSO-*d*_6_) δ ppm 4.02 (s, 3 H), 7.27 (d, *J*=9.4 Hz, 1 H), 8.01 (d, *J*=5.5 Hz, 1 H), 8.66 - 8.78 (m, 2 H), 8.83 (s, 1 H).

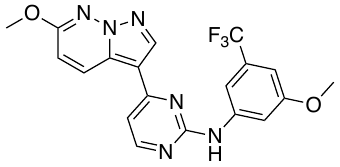

**N-(3-methoxy-5-(trifluoromethyl)phenyl)-4-(6-methoxypyrazolo[1,5-b]pyridazin-3-yl)pyrimidin-2-amine (SGC-CLK-1; CAF-170)**

3-(2-chloropyrimidin-4-yl)-6-methoxypyrazolo[1,5-b]pyridazine (0.085 g, 0.32 mmol, 1 eq.), 3-methoxy-5-(trifluoromethyl)aniline (0.075 g, 0.39 mmol, 1.20 eq.), and tert-butanol (4.5 mL) were mixed into a microwave vial, 4 small drops of TFA added, vial sealed, reaction stirred at 85 C for 15 h. Reaction mixture cooled to r.t., quenched with water and NaHCO_3_, pH adjusted to 7, solid precipitated, more water added, solid filtrated, thoroughly rinsed with water, air dried. Pale pink solid obtained, 110 mg recovered. The product was purified using Biotage Sfar 10g silica cartridge, solid load. Hexanes/EtOAc gradient from 0% to 50% EtOAc and pale whitish solid yielded (0.057 g, > 95% pure). LCMS [M+1] = 417, 418.

^1^H NMR (850 MHz, DMSO-d_6_) δ 9.90 (s, 1H), 9.02 (d, J = 9.6 Hz, 1H), 8.72 (s, 1H), 8.54 (d, J = 5.2 Hz, 1H), 7.84 (d, J = 1.8 Hz, 1H), 7.68 (t, J = 2.2 Hz, 1H), 7.43 (d, J = 5.2 Hz, 1H), 7.15 (d, J = 9.5 Hz, 1H), 6.85 (t, J = 2.0 Hz, 1H), 4.03 (s, 3H), 3.84 (s, 3H).

^13^C NMR (214 MHz, DMSO-d_6_) δ 163.22, 162.93, 162.78, 162.58, 161.21, 145.80, 141.40, 133.84, 133.54, 133.39, 132.60, 127.97, 126.70, 116.29, 113.54, 111.55, 111.00, 110.54, 106.06, 58.69, 57.91.

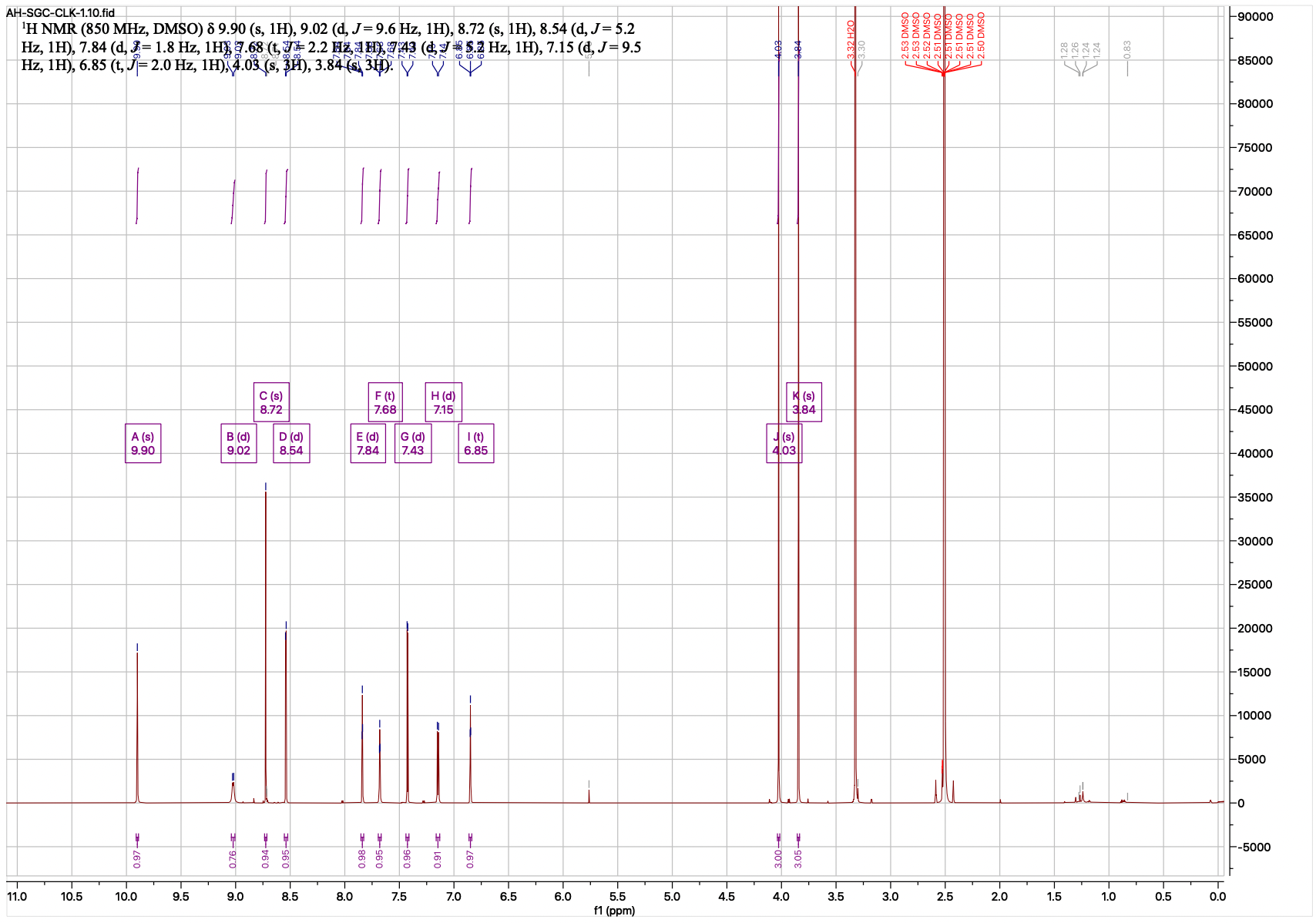

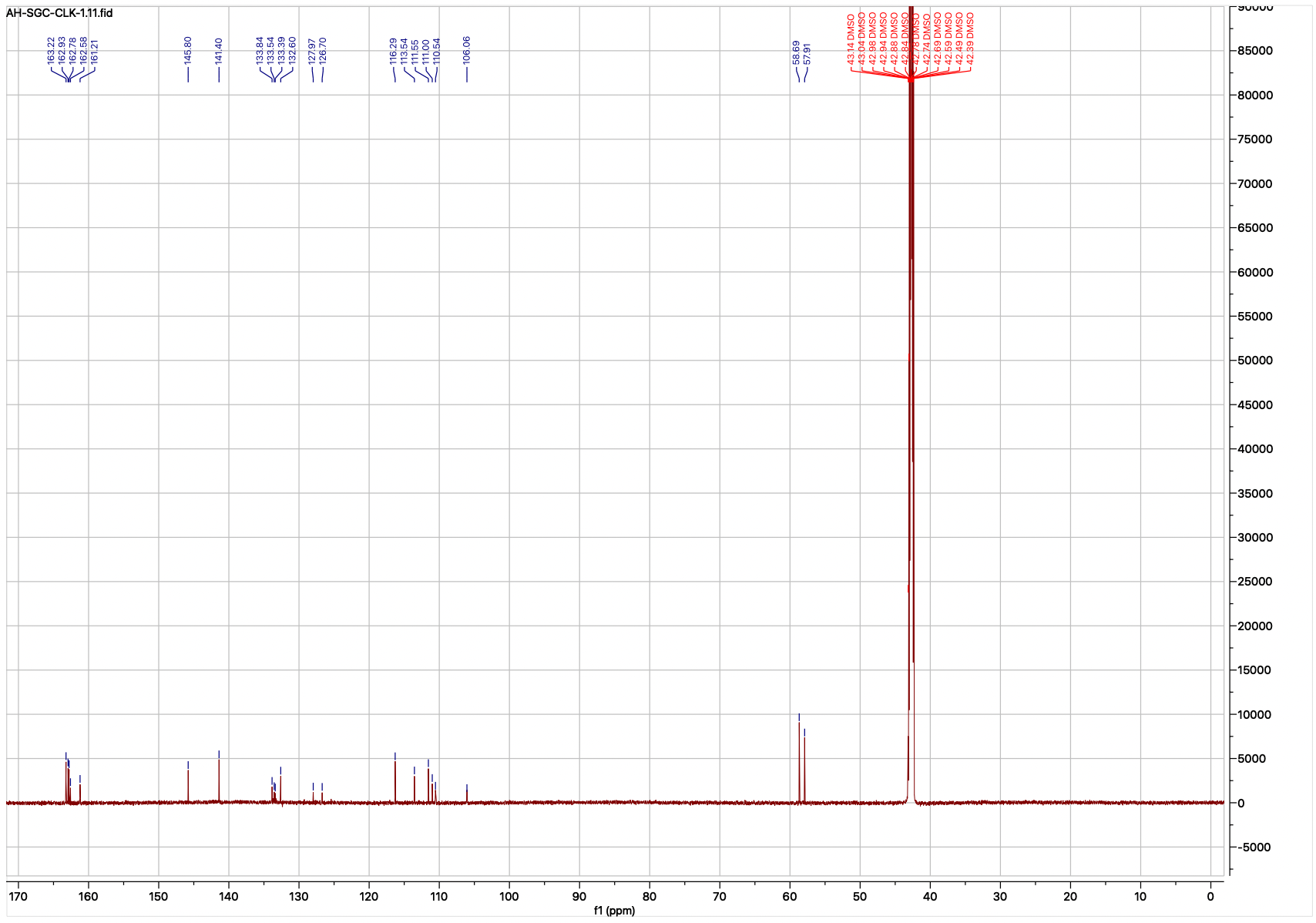

**Negative control**

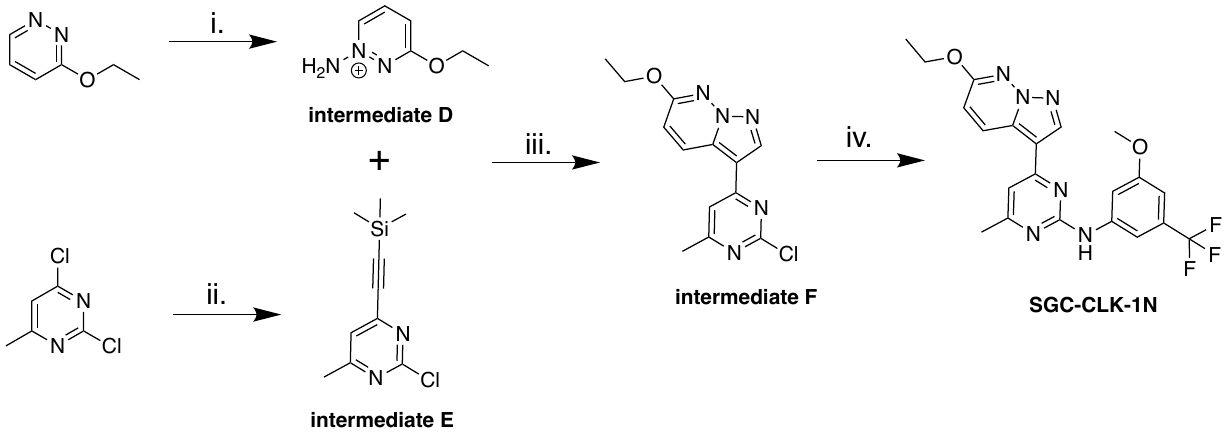

**Reagent and conditions:** i) Amino hydrogen sulphate, KHCO_3_, H_2_O, 80 C, 14 h; ii) KHCO_3_, KOH, H_2_O, DCM, r.t, 18 h; iii) ETMS, Pd(dppf)Cl_2_ · CH_2_Cl_2_, CuI, PPh_3_, TEA, THF, 70 C, 15 min; iv) TFA, tert-BuOH, 85 C, 15 h.

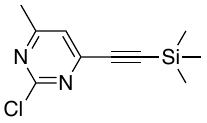

**2-chloro-4-methyl-6-((trimethylsilyl)ethynyl)pyrimidine (Intermediate E):**

2,4-dichloro-6-methylpyrimidine (1.0 g, 6.10 mmol, 1.00 eq.) and THF (30 mL) degassed with nitrogen for 10-12 min, then ethynyltrimethylsilane (0.66 g, 6.7 mmol, 1.1 eq.) and triethylamine (0.68 g, 6.7 mmol, 1.10 eq., 0.94 mL) added, degassed for another 5 min, then the rest of reagents PdCl2(dppf)-CH_2_Cl_2_adduct (0.25 g, 0.31 mmol, 0.05 eq.), copper(I) iodide (0.12 g, 0.61 mmol, 0.10 eq.), triphenylphosphine (0.16 g, 0.61 mmol, 0.10 eq.) were added. Reflux kept for 15min. Bulky solid formed. Filtered over celite, rinsed with EtOAc. Concentrated in the rotavapor, tick brown liquid/paste found. The crude product was purified using flash chromatography (0 to 15% EtOAc) 1.025 g (yield 67%), purity around 90%.

^1^H NMR (400 MHz, DMSO-*d*_6_) δ ppm 0.27 (bs, 9 H), 2.46 (s, 3 H), 7.61 (s, 1 H).

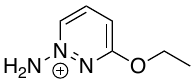

**1-amino-3-ethoxypyridazin-1-ium (Intermediate D):**

Amino hydrogen sulphate (0.82 g, 7.25 mmol, 4.5 eq.) was dissolved in water (1.6 mL), KHCO_3_ (0.72 g, 7.25 mmol, 4.5 eq.) in water (1.0 mL), pH 5. Then 3-ethoxypyridazine (0.20 g, 1.61 mmol, 1.00 eq.) was added in portion. Reaction was then stirred at 80 °C overnight. This crude is used as is for the next reaction without purification.

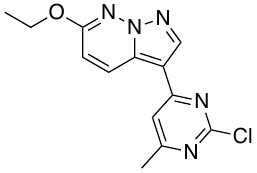

**3-(2-chloro-6-methylpyrimidin-4-yl)-6-ethoxypyrazolo[1,5-b]pyridazine (Intermediate F):**

3-ethoxy-1l4-pyridazin-1-amine (0.22 g, 1.6 mmol, 2.0 eq.) from the previous step (pH = 1) was treated with saturated KHCO_3_ to bring the pH to 7. 2-chloro-4-methyl-6-((trimethylsilyl)ethynyl)pyrimidine (0.18 g, 0.88 mmol, 1.00 eq.) was dissolved in 0.8 mL of DCM (1 M), added in one portion over the crude. KOH (0. 27 g, 4.8 mmol, 6 eq.) was dissolved in H_2_O (4.8 mL) 0.9-1.0 M and was added in one portion over previous mixture. The reaction mixture was transformed dark read in color after 5-10 min. Reaction mixture was strongly stirred at r.t. for 22 h. The crude mixture was then quenched with water, extracted with DCM, and combined organic layers dried over anhydrous Na_2_SO_4_. Crude was dry loaded on a 10 g Biotage Sfar 60 um silica cartridge (Hexanes/EtOAc 70:30) and whitish solid yielded (0.116 g, 48% yield). LCMS [M+1] = 290, purity > 95%.

^1^H NMR (400 MHz, DMSO-*d*_6_) δ ppm 1.41 (t, *J*=7.0 Hz, 3 H), 2.49 (br s, 3 H), 4.41 (q, *J*=7.0 Hz, 2 H), 7.23 (d, *J*=9.4 Hz, 1 H), 7.92 (s, 1 H), 8.69 - 8.77 (m, 2 H).

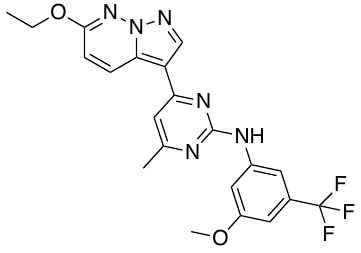

**4-(6-ethoxypyrazolo[1,5-*b*]pyridazin-3-yl)-*N*-(3-methoxy-5-(trifluoromethyl)phenyl)-6-methylpyrimidin-2-amine (SGC-CLK-1N, CAF-225):**

3-(2-chloro-6-methylpyrimidin-4-yl)-6-ethoxypyrazolo[1,5-b]pyridazine (0.065 g, 0.22 mmol, 1 eq.), 3-methoxy-5-(trifluoromethyl)aniline (0.051 g, 0.27 mmol, 1.20 eq.), and tert-butanol (4.5 mL) were mixed into a microwave vial, 4 small drops of TFA added, vial sealed, reaction stirred at 85 C for 15 h. Reaction mixture cooled to r.t., quenched with water and NaHCO_3_, pH adjusted to 7, solid precipitated, more water added, solid filtrated, thoroughly rinsed with water, air dried. Pale pink solid obtained, 110 mg recovered. The product was purified using Biotage Sfar 10g silica cartridge, solid load. Hexanes/EtOAc gradient from 0% to 50% EtOAc and pale whitish solid yielded (0.065 g, yield 30%, > 95% pure). LCMS [M+1] = 445, 446.

^1^H NMR (850 MHz, DMSO-d_6_) δ 9.85 (s, 1H), 8.98 (d, J = 9.3 Hz, 1H), 8.65 (s, 1H), 7.84 (d, J = 1.8 Hz, 1H), 7.75 (s, 1H), 7.35 (s, 1H), 7.09 (d, J = 9.5 Hz, 1H), 6.83 (t, J = 2.0 Hz, 1H), 4.42 (q, J = 7.0 Hz, 2H), 3.84 (s, 3H), 2.44 (s, 3H), 1.43 (t, J = 7.0 Hz, 3H).

^13^C NMR (214 MHz, DMSO-d_6_) δ 170.71, 163.20, 162.67, 162.40, 162.26, 145.99, 141.02, 133.78, 133.48, 133.33, 132.48, 127.99, 126.71, 116.20, 113.69, 110.82, 110.65, 110.48, 105.90, 66.43, 58.62, 27.01, 17.24.

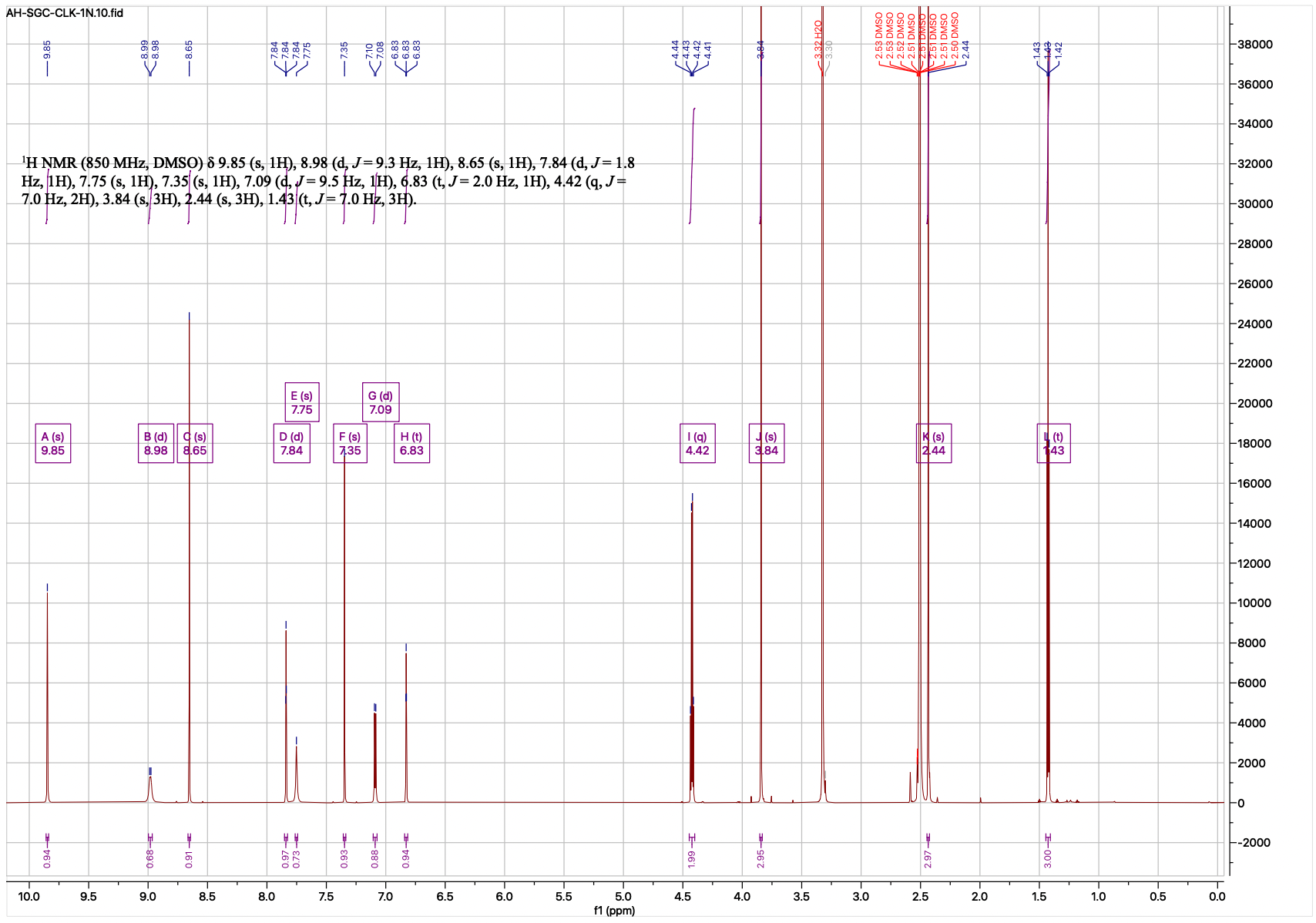

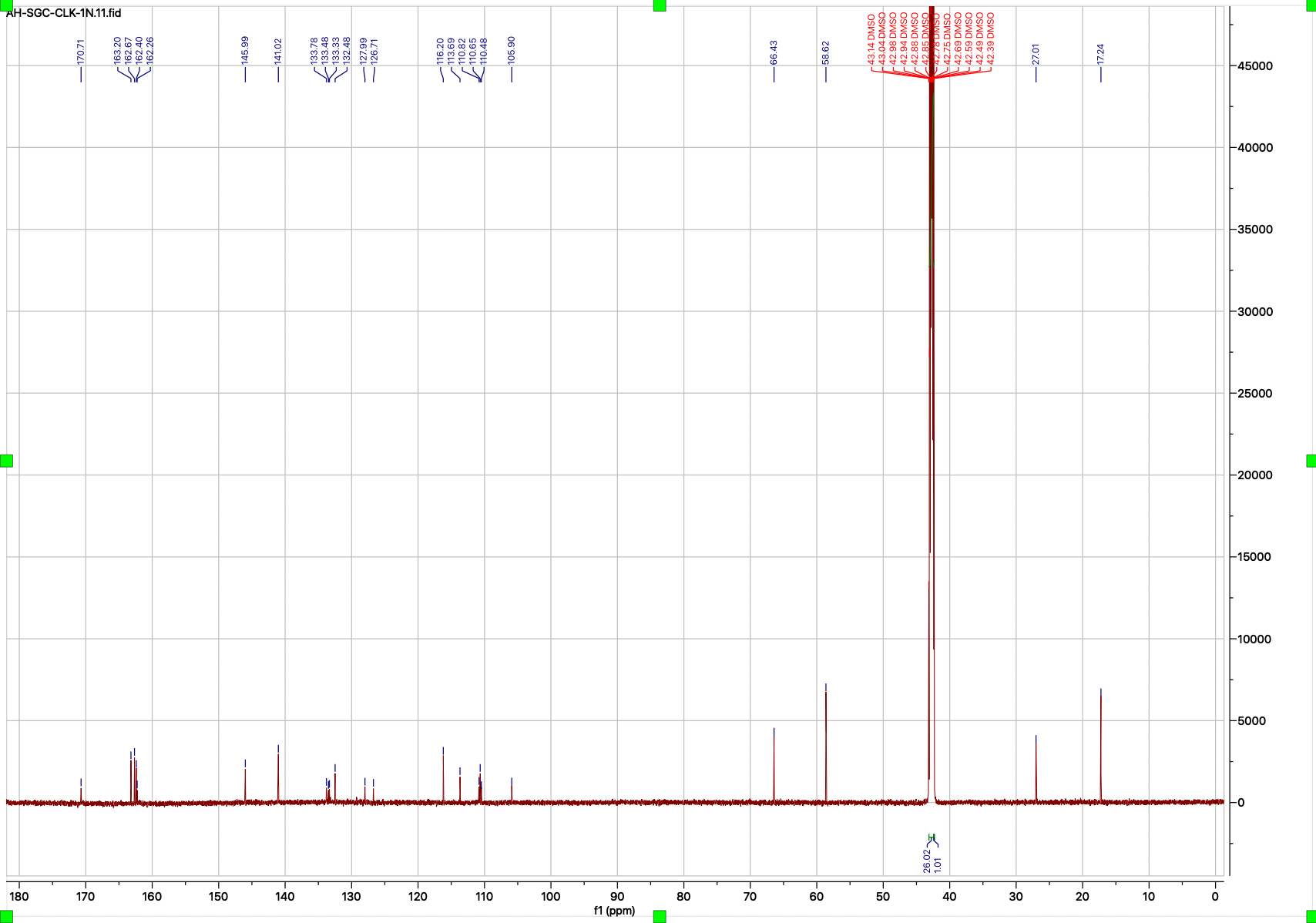
